# Comprehensive benchmarking with guidelines for analyzing transposable element-derived RNA expression

**DOI:** 10.1101/2025.09.30.679421

**Authors:** Jianqi She, Jiadong Wang, Ence Yang

## Abstract

Transposable element-derived RNAs (teRNAs) have been recognized with accelerating fundamental or pathogenic roles, especially in human. Despite the rapid development of computational methods, the best practice for accurate identification and quantification of teRNAs are currently lacking owing to the difficulties of evaluation. Here we present benchmarking of 16 representative tools with 120 simulated datasets and 60 real-world paired datasets (comprising both long- and short-read data), by evaluating the performance of teRNA identification or quantification across family-, unit-, exon-, and transcript-level. Our findings demonstrate not only the exon-level as a trade-off between accuracy and resolution for teRNA analysis, but also the level-dependent strengths and weaknesses of evaluated methods. To refine our benchmarking results, we present decision-tree-style guidelines and develop an integrated best-practice pipeline, serving as the basis for future functional researches. In addition, our evaluation framework also provides a gold standard for developing and benchmarking better computational tools in the field.

## Introduction

Transposable elements (TEs) are mobile genetic elements that constitute nearly half of the human genome [1,2]. Although most TEs are molecular fossils that have lost their transposition ability over time, accumulating evidence uncovered that numerous TEs have the ability to transcribe, producing TE-derived RNAs (teRNAs) [3–5]. These RNA molecules, including TE-only teRNAs (autonomous transcription of TE units) and TE-gene chimeric RNAs (co-transcription with genes), have been increasingly recognized for their roles in embryo development [6], immune response [7], aging [8] as well as complex diseases, such as cancer [9,10] and neurodegenerative diseases [11]. For example, HERVH-derived RNAs interact with pluripotency-associated transcription factors to regulate the expression of neighboring genes and maintain human embryonic stem cell identity [12]. Moreover, recent studies have also revealed that teRNAs could predict the outcome of immune checkpoint blockade in human lung adenocarcinoma and hold potential as dominant anti-tumor antibody targets [13]. Given diverse physiological and pathological roles acknowledged, accurate detection and quantification of teRNAs has become an urgent issue for identifying functional teRNAs.

With the rapid advancement of next-generation sequencing technology, various methods have been developed to identify and quantify teRNA expression from Illumina paired-end short reads [14]. However, these methods differ significantly in the research levels of teRNAs. Given the repetitive nature of TEs, most studies focus on family-level by combining TE expression within certain family to avoid ambiguously alignment, while the others have explored approaches at more granular levels, concentrating on individual TE unit (unit-level), TE-derived exons (exon-level) or TE-derived transcripts (transcript-level), to detect functional teRNAs at specific genomic locations. For unit-level and exon-level, we define a TE unit as a specific genomic locus occupied by a TE, while a TE exon denotes a transcript segment derived from such a locus (A glossary of key terms on teRNAs is provided in Table S1 for reference). Thus, for a TE-only transcript that is derived from a single genomic TE locus (e.g., a transcribed L1 element), the unit, exon, and transcript are one and the same. Additionally, even within the same analytical level, different tools usually yield varying results, which largely limits the capture of functional teRNAs [15]. Therefore, given the diversity and difference of teRNA analysis tools, a reliable and systematic benchmarking study is necessary to accelerate teRNA studies.

The lack of a ground-truth annotation makes it challenging to systematically analyze the accuracy of teRNA detection and quantification tools across different teRNA levels. In this study, we address this gap by establishing essential ground truths for teRNAs using ultra-deep long-read technology and generating 180 benchmarking datasets (including 120 simulated datasets and 60 independent real-world datasets) to evaluate the performance of 16 teRNA analysis tools across family-, unit-, exon-, and transcript-levels. We identified not only the level-dependent strengths and weaknesses of evaluated methods, but also a best-practice pipeline, termed TERA, which achieved the best performance in both teRNA detection and quantification. Finally, by applying TERA to RNA-seq datasets from glioblastoma patients, we successfully conducted a thorough analysis of teRNA expression profiles and pinpointed functional teRNAs with oncogenic potential. Overall, our study offers practical guidance for detecting and quantifying teRNA expression from next-generation sequencing data, which will speed up the exploration of functional teRNAs.

## Results

### Benchmarking framework based on full-length transcriptome

To rigorously evaluate and compare the existing methods for detecting TE expression, we constructed ground truths for teRNAs based on the full-length transcriptome from HEK293T and U251 cells (Figure 1A). By performing ultra-deep long-read (PacBio) and short-read (Illumina) RNA sequencing, we obtained 66,397 and 112,313 high-quality full-length teRNA transcripts (each containing at least one TE unit) in HEK293T and U251 cells, respectively. Here, high-quality teRNAs were defined by a stringent set of criteria: (1) for multi-exon transcripts, the supporting HiFi reads should be ≥ 2 or the splicing junction should be supported by ≥ 1 junction read from short-read RNA-seq data or transcript annotation; (2) for mono-exon transcripts, supporting HiFi reads should be ≥ 2; (3) sequence length of TEs within teRNAs should be ≥ 20 bp. Compared with the GENCODE-annotated transcripts, we identified 22,225 (33.47%) novel teRNAs in HEK293T and 47,744 (42.5%) in U251. Most novel teRNAs are TE-gene chimeric transcripts (chimeric teRNAs; HEK293T: *n* = 20,216; U251: *n* = 44,655), while the others are TE-only teRNAs, which are further categorized into intronic teRNAs (HEK293T: *n* = 1048; U251: *n* = 1320), intergenic teRNAs (HEK293T: *n* = 468; U251: *n* = 574) and antisense teRNAs (HEK293T: *n* = 493; U251: *n* = 1195) based on their locations relative to annotated genes (Figure 1B, Figure S1A).

**Figure 1.**
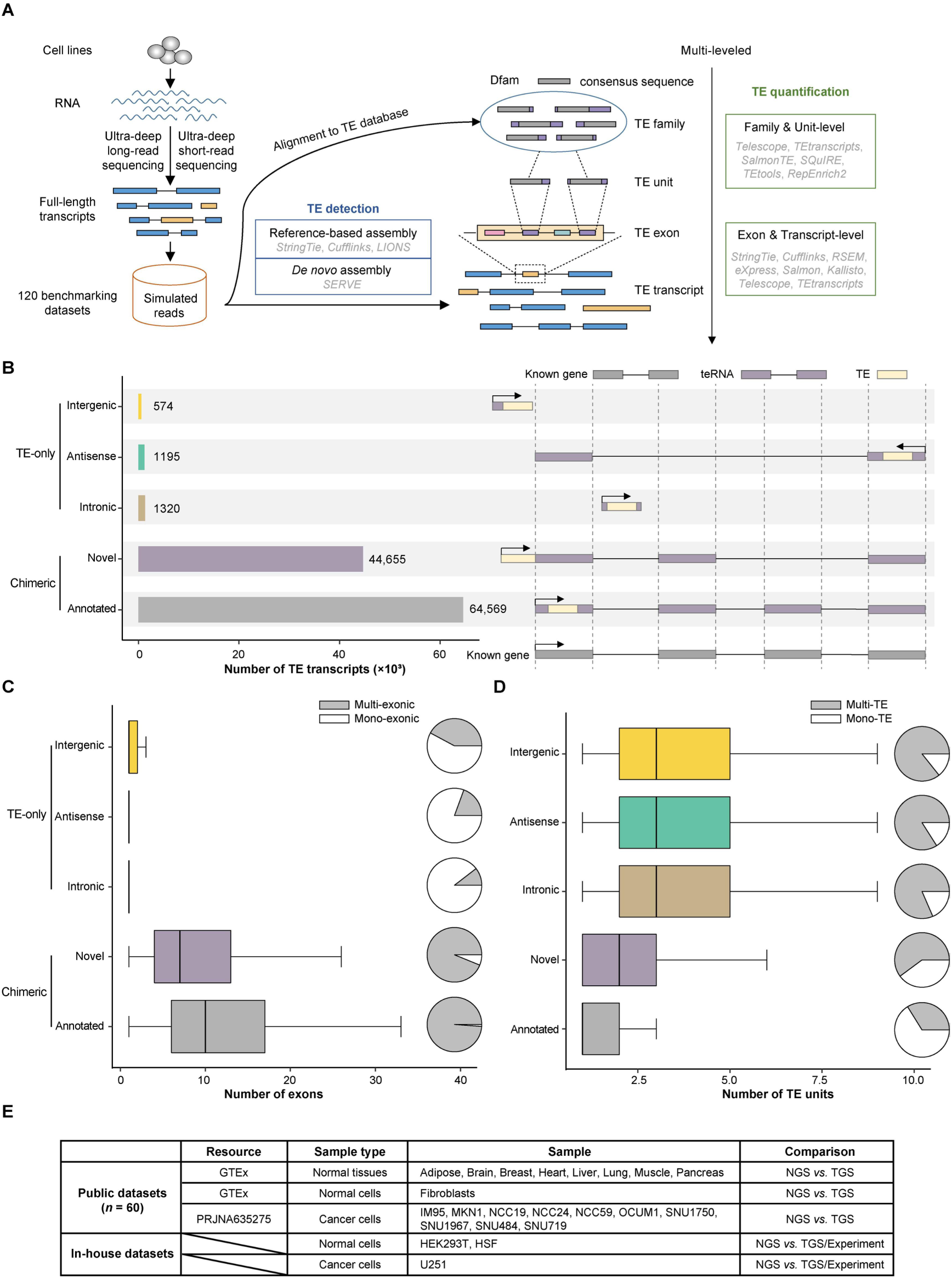
Overview of benchmarking framework. **A**. Schematic overview of the experimental design and benchmarking framework for teRNA detection and quantification methods. **B**. The number of full-length chimeric teRNAs (annotated and novel) and TE-only teRNAs (intergenic, antisense and intronic) detected from U251 cells. **C–D**. The number of exons (**C**) and TE units (**D**) of teRNAs stratified by their transcript classification in U251 cells. The center line indicates the median, the limits are the interquartile range (IQR), the whiskers represent 1.5× the IQR. **E**. Summary of real-world datasets for benchmarking teRNA detection and quantification methods.

Novel chimeric teRNAs have significantly fewer exons than annotated teRNAs but more than TE-only teRNAs (Wilcoxon rank-sum test, *p*-value < 0.001), most of which are mono-exonic (particularly intronic teRNAs) (Figure 1C, Figure S1B). In contrast, most of chimeric teRNAs contain only one TE unit, significantly fewer than TE-only teRNAs (Wilcoxon rank-sum test, *p*-value < 0.001), which are often “tandem-TE” transcripts (Figure 1D, Figure S1C). In both cells, less than 1% of teRNA transcripts originate from a single TE locus (Figure S2A), which suggests that most TEs tend to co-transcribe with nearby genes or TEs rather than transcribing independently as mono-TE. We further characterized the TE units composing these exons and found that mammalian-wide interspersed repeat (MIR), Alu, and L2 families were significantly enriched in TE-gene chimeric exons (*p*-value < 0.001; Figure S3; Table S2). Notably, for TE-only exons, L1 families (particularly L1HS_5end and L1P1_orf2) were significantly enriched in HEK293T cells, while the HERVH family was significantly enriched in U251 (*p*-value < 0.001; Figure S3; Table S2). Furthermore, about half of all TE exons contained more than one TE unit (Figure S2B), and specific TE combinations, such as L1HS_5end:L1P1_orf2 in HEK293T and HERVH:LTR7Y in U251, were significantly co-enriched within individual exons (permutation test *p*-value < 0.001; Figure S4; Table S3), revealing complex patterns of TE exon formation. Additionally, about half of chimeric teRNAs contain TE transcription start sites (TE-TSS teRNAs, enriched in the ERV family), TE splicing sites (TE-SS teRNAs, enriched in the SVA family), or TE transcription termination sites (TE-TTS teRNAs, enriched in the Alu family), indicating the complicated regulatory potential of TEs in the genome (Figure S5).

Given the complexity and diversity of teRNAs, we constructed 120 benchmarking datasets by simulating short-read RNA sequencing data with various sequencing depths (10–100 million paired-end (PE) reads) and read lengths (50–300 bp) based on full-length transcriptome from HEK293T and U251 as the ground truth (Figure 1A). For evaluation, we selected 4 teRNA detection tools (StringTie [16], Cufflinks [17], LIONS [18], and SERVE [19]) and 12 teRNA quantification tools (Telescope [20], TEtranscripts [21], TEtools [22], SQuIRE [23], SalmonTE [24], RepEnrich [25], StringTie, Cufflinks, RSEM [26], eXpress [27], Salmon [28], and Kallisto [29]). Among the detection tools, reference-based assembly methods (StringTie, Cufflinks, and LIONS) aim to identify TE-gene chimeric transcripts (chimeric teRNAs) while SERVE, based on *de novo* assembly, is designed to detect TE-only teRNAs. teRNA quantification tools differ in the handling of multi-mapping reads and the research levels of teRNAs. A summary of the input, output, and working principles for all benchmarked tools is provided in Table S4. Thus, we systematically evaluated and compared the accuracy of teRNA detection and quantification across family-, unit-, exon-, and transcript-levels (Figure 1A). In addition to simulated benchmarking datasets, we also introduced 60 public datasets from normal tissues [30], normal cells [19,30] and cancer cells [31], with each sample include both long-read and short-read RNA-seq data for method evaluation (Figure 1E).

### Evaluation of teRNA detection tools

By applying the teRNA detection tools to the simulated benchmarking datasets, we observed that all tools achieved high F_1_ scores at the family-level but showed a substantial drop at more granular levels, particular at the transcript-level (< 0.3 in all datasets), indicating the challenge of accurate locus-specific teRNA identification (Figure 2A and B, Figure S6A and B; Table S5). The performance of all detection tools improved with increasing sequencing depth and read length across all levels. For family-level, StringTie (guided) is the most sensitive (average recall = 0.923) and accurate (average F_1_ score = 0.949) across nearly all datasets (Figure 2E, Figure S6E; Table S5). In contrast, SERVE achieved significantly higher F_1_ scores (average F_1_ scores = 0.668) than the other tools (average F_1_ scores < 0.4) for detecting teRNA at the unit-level in case that read length was PE150 or longer (Wilcoxon rank-sum test, *p*-value < 0.001). However, SERVE was more sensitive to changes in sequencing depth and read length, and StringTie (guided) would be a better alternative for datasets with shorter read length (< PE150) and lower sequencing depth (< 50 million) for unit-level.

**Figure 2.**
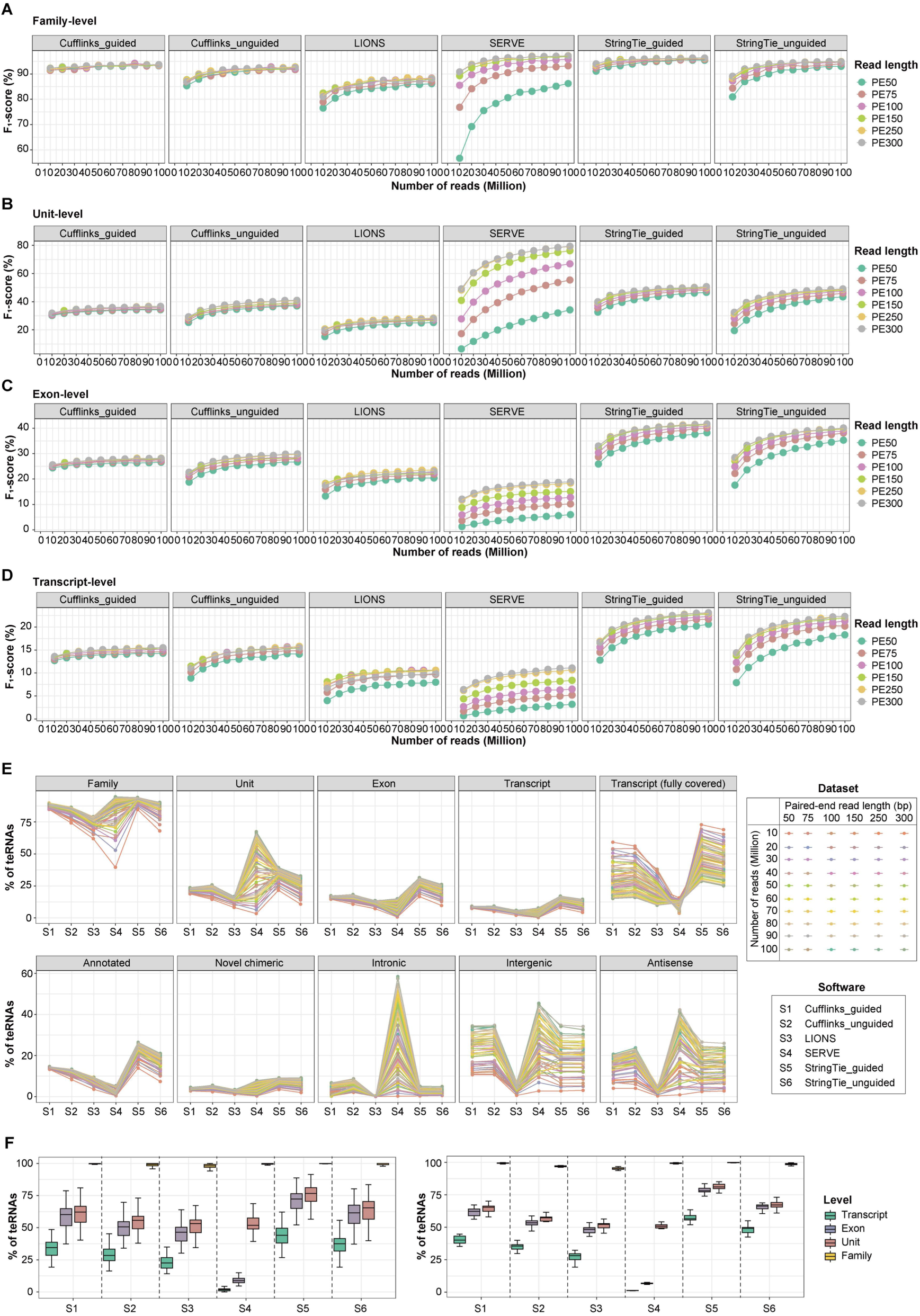
Evaluation of teRNA detection methods across different levels. **A–D**. The performance of teRNA detection methods at the family- (**A**), unit- (**B**), exon- (**C**) and transcript-levels (**D**) in simulated datasets from U251 cells. **E**. Sensitivity of teRNA detection methods in the identification of different levels and types of teRNAs in simulated datasets from U251 cells. **F**. The performance of teRNA detection methods across different levels in real-world datasets from normal tissues/cells (left) and cancer cells (right). The center line indicates the median, the limits are IQR, the whiskers represent 1.5× the IQR.

For transcript-level detection, StringTie (guided) detected the highest number of chimeric teRNAs, while SERVE detected the highest number of TE-only teRNAs, particularly intronic teRNAs (Figure 2E, Figure S6E). However, none of the examined methods identified more than 25% of teRNA transcripts in any simulated datasets, which largely limited the discovery of functional teRNAs (Figure 2D, Figure S6D; Table S5). We also assessed the ability of each tool to discover novel teRNAs (those not present in reference annotations). The detection of novel teRNAs proved exceptionally challenging, with sensitivity rates for all tools below 5% across almost all datasets (Figure S7A). Among the teRNAs that each tool successfully detected, SERVE demonstrated a superior ability to discover novel teRNAs, which constituted up to 73% of its total predictions. In contrast, the proportion of novel teRNAs detected by other tools was consistently below 30% (Figure S7B). Additionally, the proportion of novel teRNAs discovered by all tools increased with sequencing depth, indicating that deeper sequencing is a key strategy to uncover new functional teRNAs.

Although exon-level detection is less commonly studied, our results showed that the F_1_ scores for teRNA-exons were significantly higher than for teRNA-transcripts in all datasets (Wilcoxon rank-sum test, *p*-value < 0.001; Figure 2C, Figure S6C; Table S5). StringTie (guided) was the most accurate at the exon-level, with F_1_ scores reaching 0.486, which is twice higher than the score at transcript-level. In the 60 independent real-world datasets, the sensitivities of all the tools at the exon-level were also significantly higher than the transcript-level (Wilcoxon rank-sum test, *p*-value < 0.001) and close to the unit-level (Figure 2F). Therefore, with higher resolution than unit-level and greater accuracy than transcript-level, the exon-level may be a better alternative for detecting teRNAs.

In addition to read length and sequencing depth, teRNA expression levels also influenced the accuracy of teRNA detection. Our results showed that all software achieved significantly higher sensitivity in the detection of highly expressed (Figure S8) and reads-fully-covered teRNA-transcripts (Wilcoxon rank-sum test, *p*-value < 0.001; Figure 2E, Figure S6E). To further illustrate the influence of teRNA expression, we simulated datasets with varying teRNA expression levels: lowly-expressed (LTE: FPKM > 0), medium-expressed (MTE1: FPKM > 0.05; MTE2: FPKM > 0.1), and highly-expressed (HTE: FPKM > 1). As expression levels increased from LTE to HTE, F_1_ scores and recall improved significantly, while precision remained relatively stable across all levels for all tools (Figure 4G, Figure S10A). Notably, at both the transcript and exon levels, the recall for TE-only teRNAs increased more than for chimeric teRNAs, particularly with SERVE (Figure 3, Figure S9).

**Figure 3.**
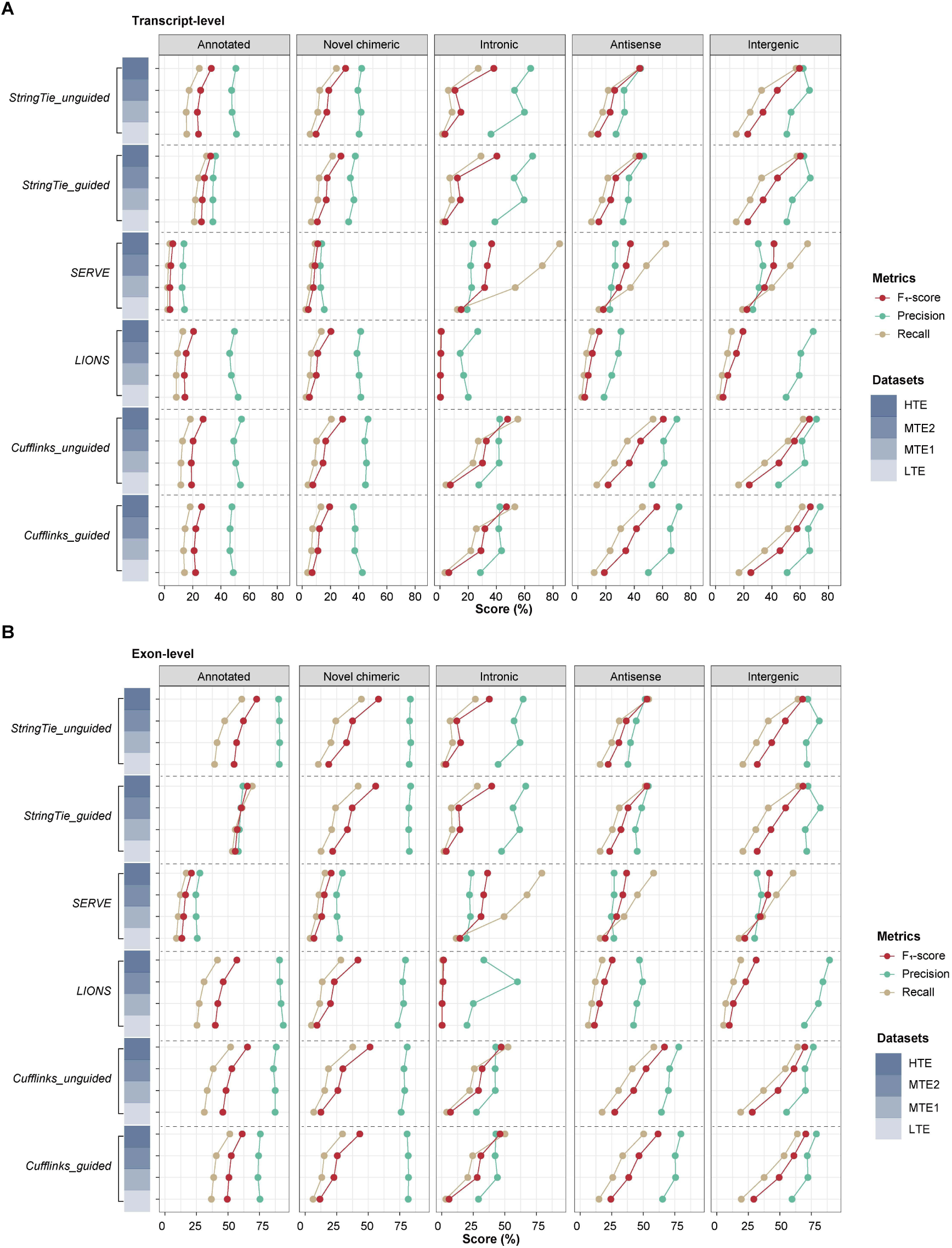
Performance of methods in the detection of teRNAs with different expression level. **A–B**. The performance of teRNA detection methods in the identification of different types of teRNA-transcripts (**A**) teRNA-exons (**B**) in LTE, MTE1, MTE2 and THE datasets from U251 cells.

**Figure 4.**
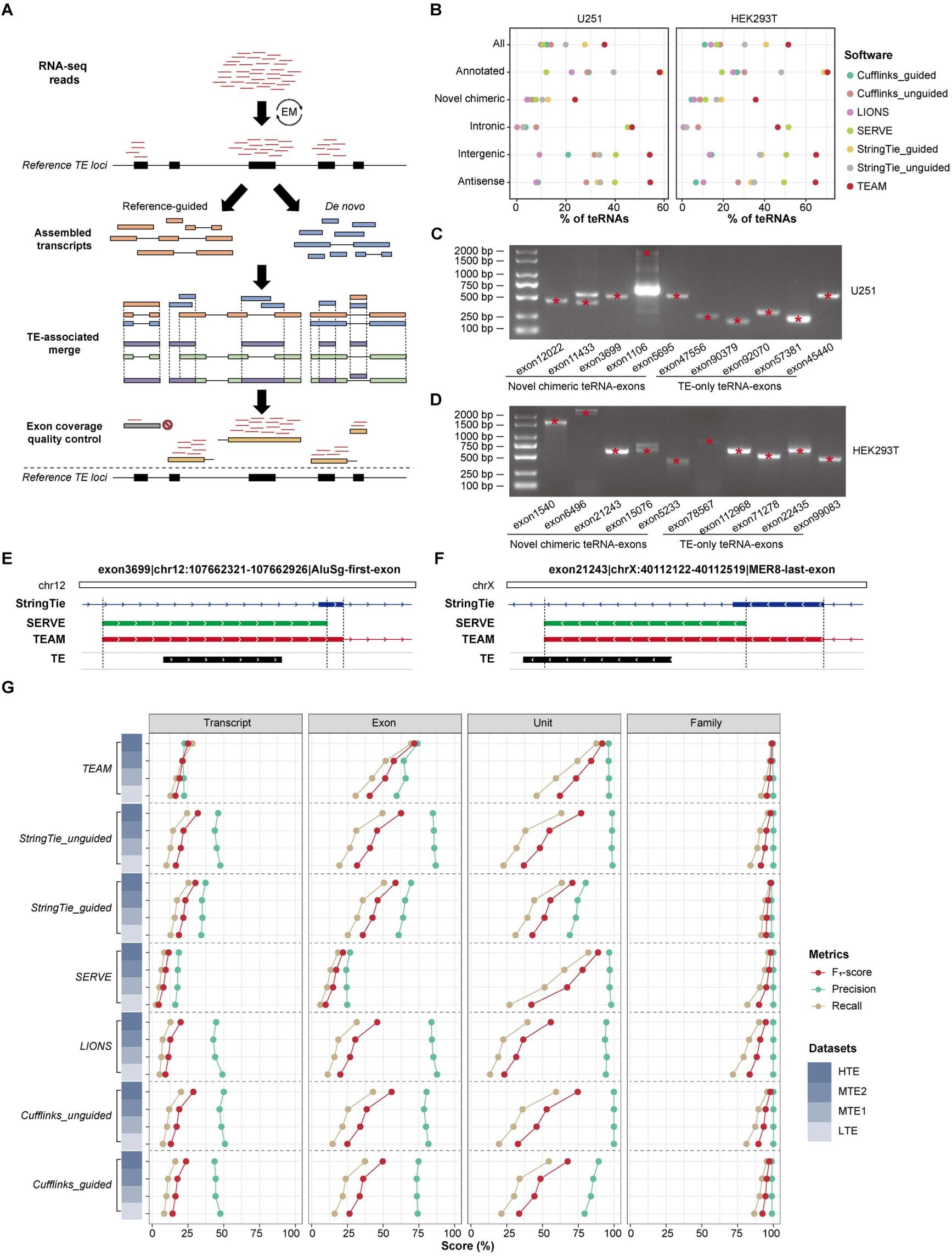
The performance of TEAM in teRNA detection. **A**. The scheme of TEAM. **B**. The sensitivity of TEAM in the identification of different types of teRNA-exons in RNA-seq data from U251 (left) and HEK293T (right) cells. **C–D**. The experimental validation of TEAM-identified teRNA-exons in U251 (**C**) and HEK293T (**D**) cells. Red star points indicate the target band. **E–F**. The example of TEAM-identified teRNA-exons: AluSg-first-exon (**E**) and MER8-last-exon (**F**). **G**. The performance of TEAM and existing teRNA detection methods across different levels in simulated datasets from U251 cells.

### Integration of teRNA detection tools achieves better accuracy

Given StringTie’s effectiveness on chimeric teRNAs and SERVE’s sensitivity in TE-only teRNAs, we developed an integrated algorithm (TEAM) that combines these tools and focuses on detecting teRNA-exons as well as teRNA-units and teRNA-families (Figure 4A). To maximize detection sensitivity and maintain precision, TEAM integrates transcript assembly results of StringTie and SERVE, following a two-step read assignment for quality control (see details in Materials and methods). Especially for detecting chimeric teRNA-exons, TEAM leverages the sensitivity of SERVE in teRNA-units to refine the boundaries of StringTie-identified teRNA-exons, contributing to more accurate detection of chimeric teRNA-exons (Figure 4E and F).

By applying TEAM to the benchmarking simulated datasets, we showed that TEAM achieved the best performance in detecting teRNA-exons in almost all datasets (Wilcoxon rank-sum test, *p*-value < 0.001; Figure 4G, Figure S10A). Similar to SERVE, the accuracy and sensitivity of TEAM was improved with increasing sequencing depth and read length (Figure S11). Additionally, the accuracy and sensitivity of TEAM in detecting teRNA-exons substantially increased from LTE to HTE datasets (Figure 4G, Figure S10A). Particularly in HTE datasets, TEAM achieved the best trade-off between precision and recall and the F_1_ scores reached over 70% (0.718 in U251 and 0.777 in HEK293T). Besides exon-level, TEAM was also the most accurate and sensitive tool at both the unit-level and family-level in almost all the simulated datasets, while accurately detecting teRNA-transcripts still remained a challenge for TEAM, similar to other tools (Figure 4G, Figure S10A).

To further evaluate the performance of TEAM in real-world datasets, we firstly applied TEAM to in-house RNA-seq data from HEK293T and U251 cells, and confirmed that TEAM still achieved the highest sensitivity in detecting both chimeric and TE-only teRNAs (Figure 4B). To assess the precision of TEAM, we randomly selected 10 teRNA-exons (5 chimeric teRNA-exons and 5 TE-only teRNA-exons) that detected solely by TEAM from each cell line for experimental validation (Figure 4C and D). In both U251 and HEK293T cells, all the 20 teRNA-exons exclusively identified by TEAM were verified, demonstrating TEAM’s high precision in detecting teRNA-exons. In the 60 independent real-world datasets, TEAM consistently demonstrated high sensitivity across all levels of teRNA detection (Figure S10B).

### Evaluation of teRNA quantification tools

To evaluate the performance in teRNA quantification, we applied 12 teRNA quantification tools to the 120 benchmarking simulated datasets and calculated the Pearson correlation coefficient between the numbers of predicted and simulated teRNA reads across all levels. As expected, quantification of teRNA at the family-level were more accurate than locus-specific levels (Figure 5A, Figure S12A). Among the tools, Telescope was the most accurate and robust in quantification at both the family-level and unit-level. However, it still remains challenging for all the examined methods (particularly SalmonTE) to accurately quantify younger teRNA-families (defined by a lower Kimura divergence score), primarily due to the high sequence identity among internal members, though these methods develop algorithms to improve the accuracy of TE-reads assignment (Figure 5B, Figures S12B and S13A). For instance, HERV9NC, a very young family (Kimura < 5), showed poor accuracy in locus-specific quantification (Figure S13B), while the quantification of old HERVL40 family (Kimura > 25) is highly accurate across all methods (Figure S13C).

**Figure 5.**
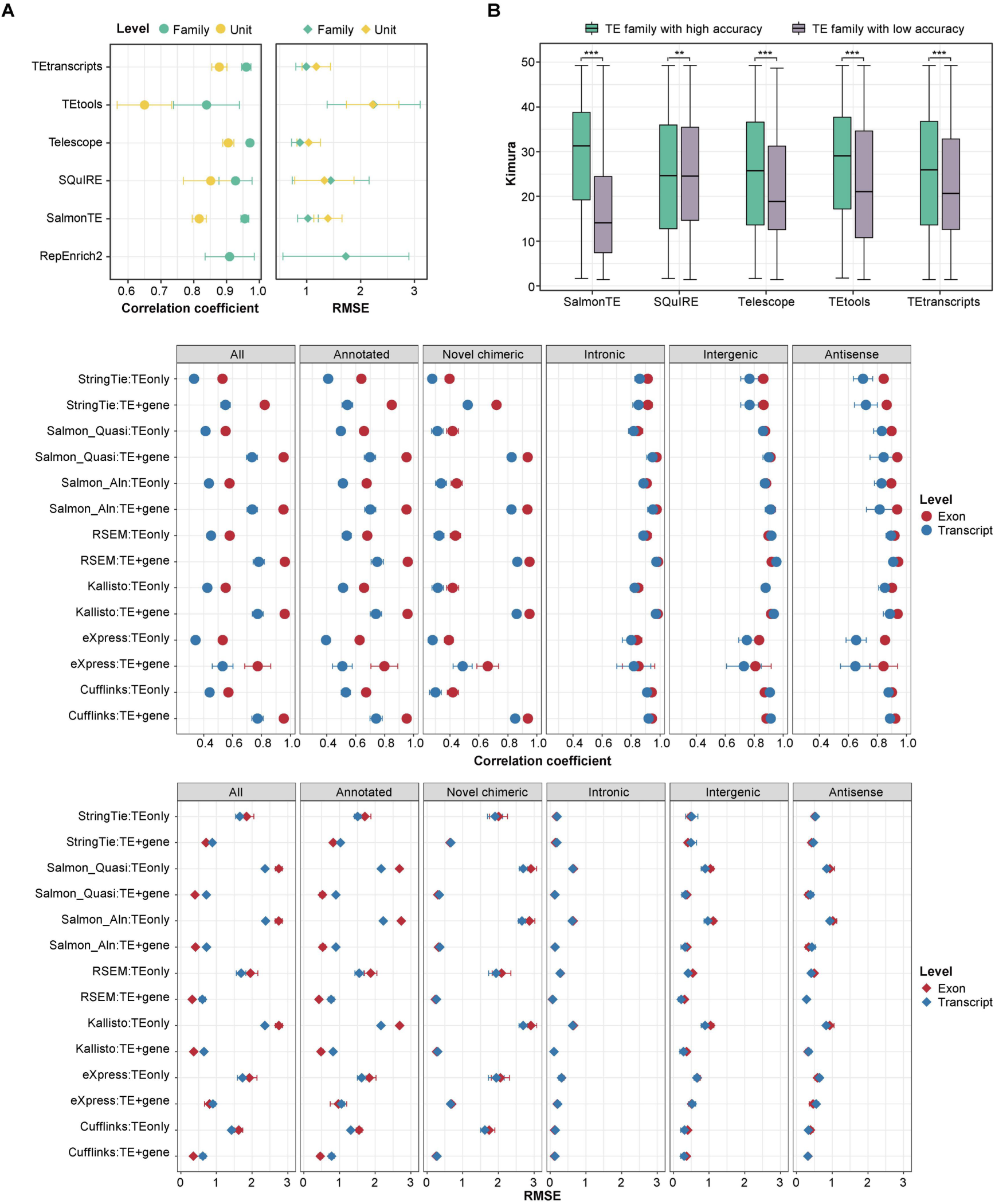
Evaluation of teRNA quantification methods. **A**. The performance of teRNA quantification methods at the teRNA-unit and teRNA-family level. **B**. Kimura’s distance of teRNA-families with high and low quantification accuracy in U251 cells. The center line indicates the median, the limits are IQR, the whiskers represent 1.5× the IQR. *, *p*-value < 0.05; **, *p*-value < 0.01; ***, *p*-value < 0.001. **C**. The performance of teRNA quantification methods at the transcript-level and exon-level.

For teRNA-transcripts, all the software offer two quantification strategies: TEonly (TEO) and TE+gene (TEG), depending on the reference transcripts include teRNAs only or appended with the other genes. We found that all the methods in both strategies performed well in the quantification of TE-only teRNAs at transcript level, as most of TE-only teRNAs were mono-exonic (Figure 5C, Figure S12C). In contrast, the accuracy of chimeric teRNA-transcript quantification using TEO strategies was obviously lower than TEG strategies, due to the impact of isoform quantification. Among these software, RSEM (average 0.781), Kallisto (average 0.774) and Cufflinks (average 0.773) achieved better performance in the quantification of all teRNA-transcripts.

For teRNA-exons, we normalized teRNA-transcript expression level by exon length. Similarly, all methods in both TEG and TEO strategies achieve high performance in TE-only teRNA-exon quantification, while TEG strategies outperformed TEO strategies in chimeric teRNA-exon quantification (Figure 5C, Figure S12C). Compared to the teRNA-transcript level, the accuracy of teRNA-exon quantification was significantly higher, especially for chimeric teRNAs (Wilcoxon rank-sum test, *p*-value < 0.001). Among the examined software, the correlation coefficients of RSEM, Kallisto and Salmon (both alignment-based and quasi-mapping-based) reached over 0.95. To further compare the quantification performance of teRNA-exons and teRNA-units, we introduced human fibroblasts datasets and randomly selected 5 teRNAs for experimental validation. The results indicated that teRNA-exon quantification (particularly by Cufflinks, RSEM and alignment-based Salmon) achieved better performance than teRNA-unit quantification (Figure S14).

### Best practice of teRNA analysis pipeline

Based on our evaluation, we provided level-specific recommendation and guidelines for users in both detection and quantification of teRNAs (Figure 6A). TEAM (the integration of reference-based assembly and *de novo* assembly) proved effective in most scenarios for teRNA detection. To simplify the analysis process, we also developed a user-friendly best-practice pipeline, TERA, suitable for teRNA analysis at the family-, unit- and exon-level in most datasets (Figure 6B). TERA includes one detection mode, one annotation mode and two quantification modes: (1) detection mode utilizes StringTie and SERVE for teRNA assembly and TEAM for the detection and quality control of teRNA-exons; (2) annotation mode annotate teRNAs with TE annotation from Dfam database and host gene annotation from GENCODE; (3) quantification mode 1 uses Telescope for the quantification of teRNA-loci and teRNA-families; (4) quantification mode 2 employs RSEM or Kallisto for the quantification of teRNA-exons. Unlike existing integrated pipelines such as SQuIRE, TEToolkit, and TEtools, which primarily focus on unit- or family-level quantification, TERA provides comprehensive detection and quantification across transcript, exon, unit, and family levels, with superior accuracy in overlapping functionalities (Table S6).

**Figure 6.**
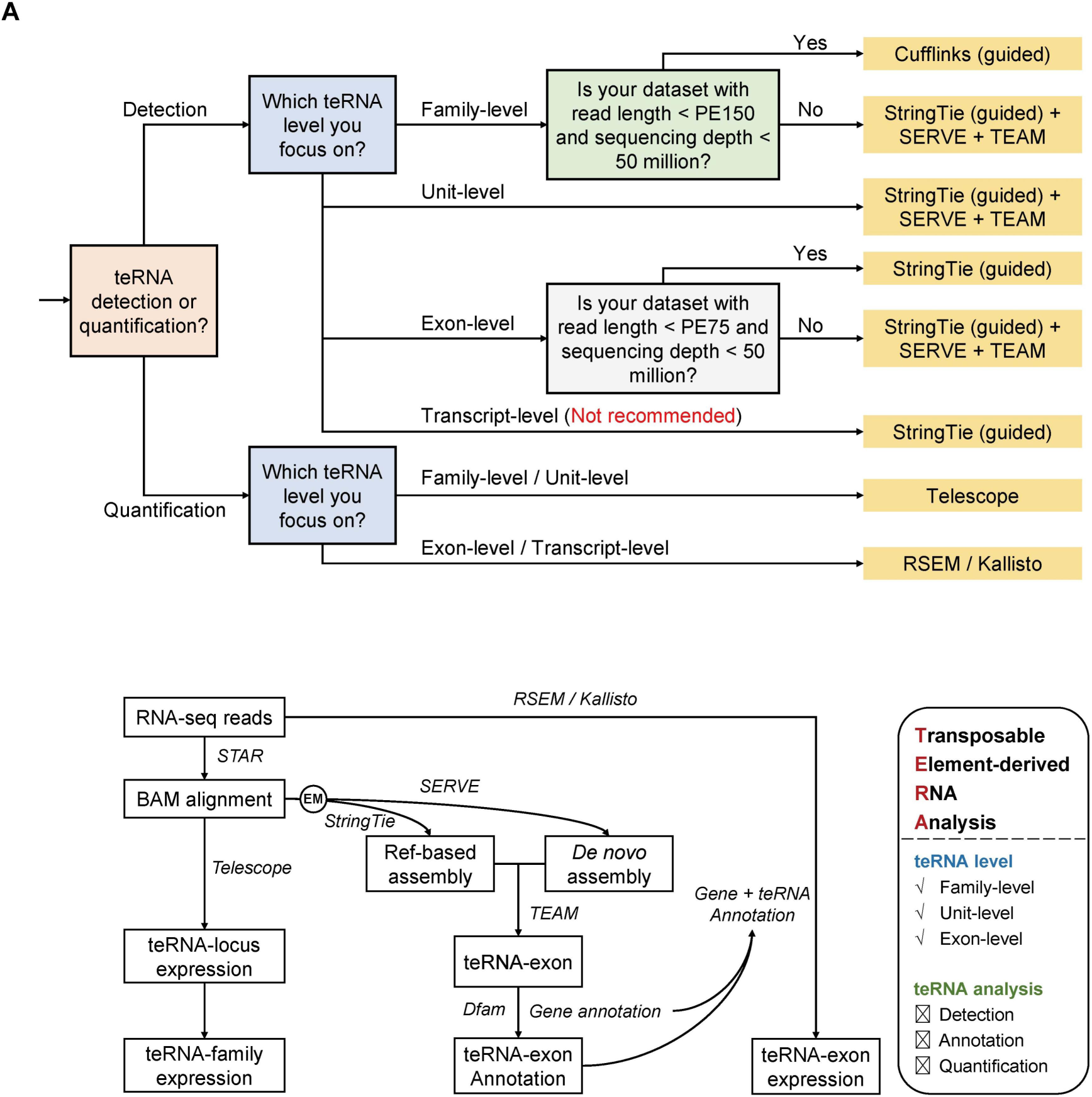
teRNA analysis guidelines for users. **A**. Practical guidelines for users in teRNA analysis across four levels. **B**. Workflow of TERA in teRNA detection and quantification.

To demonstrate the utility of TERA in analyzing teRNA expression profiles and identifying functional teRNAs, we applied the pipeline to RNA-seq data of normal and tumor brain samples from 100 glioblastoma patients [32]. In total, 74,150 teRNA-exons, 133,694 teRNA-units and 1134 teRNA-families were detected and quantified in these samples. We detected a total of 60,709 teRNAs in the tumor samples compared to 48,536 in the normal sample, suggesting a substantial reactivation of TEs in glioblastoma. Through dimensionality reduction using gene expression level as well as teRNA-exon, -unit, and -family expression levels, our results revealed that the tumor and normal samples were clearly distinguished using genes (silhouette coefficient = 0.59) and teRNA-exons (silhouette coefficient = 0.53), and to a lesser extent by teRNA-families (silhouette coefficient = 0.45) and teRNA-units (silhouette coefficient = 0.32) (Figure 7A), suggesting that teRNA-exons can also be served as potential biomarkers for distinguishing tumor from normal tissue samples.

**Figure 7.**
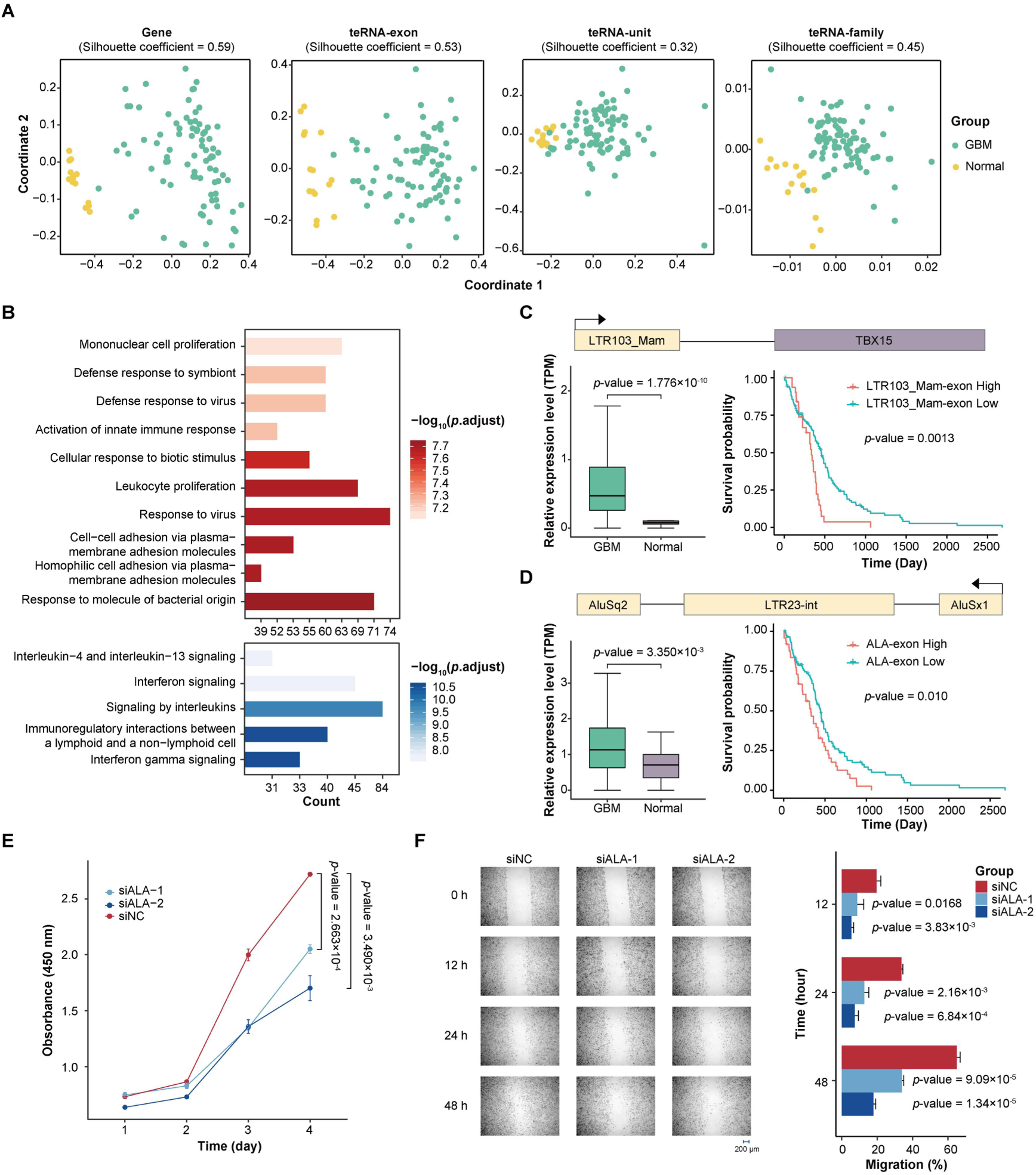
Differentially expressed teRNAs in glioblastoma. A. Multi-dimensional analysis using gene expression level as well as teRNA-exon, -unit, and -family expression levels. **B**. Gene ontology (top) and Reactome pathway (bottom) enrichment analysis of glioblastoma-upregulated chimeric teRNA-exons. **C**. LTR103_Mam-exon expression in glioblastoma and normal brain tissues (left) and Kaplan-Meier analysis of overall survival in glioblastoma patients from TCGA datasets (right). **D**. ALA-exon expression in glioblastoma and normal brain tissues (left) and Kaplan-Meier analysis of overall survival in glioblastoma patients from TCGA datasets (right). **E**. The effect of ALA knockdown on cell growth rate as determined by CCK-8 assay in U251 cells. **F**. The effect of ALA knockdown on cell migration in U251 cells as measured by scratch migration assay. The center line indicates the median, the limits are IQR, the whiskers represent 1.5× the IQR.

To further explore the potential role of teRNA-exons in glioblastoma, we conducted differential expression (DE) analysis and identified 13,041 upregulated teRNA-exons, including 6221 chimeric teRNA-exons and 6820 TE-only teRNA-exons (Table S7). Gene ontology and pathway enrichment analyses of the upregulated chimeric teRNA-exons highlighted significant involvement in immune response activation (e.g., interferon signaling) and cell adhesion processes (Figure 7B). Additionally, by integrating RNA-seq data with clinical information from The Cancer Genome Atlas (TCGA) [33] and Chinese Glioma Genome Atlas (CGGA) [34] datasets, we identified 141 glioblastoma-upregulated chimeric teRNA-exons significantly associated with worse prognosis in glioblastoma patients in both datasets (*p*-value < 0.05; Table S8). Among these teRNA-exons, LTR103_Mam-exon, the first exon of TBX15 gene, has been reported as a malignant accelerator in various cancers (Figure 7C, Figure S15A).

Besides chimeric teRNA-exons, we also assessed the impact of TE-only teRNA-exons on glioblastoma. Considering that teRNA expression may be influenced by host genes and nearby genes, we calculated the independent expression score of teRNA-exons and identified 8 independent TE-only teRNA-exons significantly upregulated and associated with poor prognosis in glioblastoma (Table S9). Among these teRNA-exons, AluSq2–LTR23-int–AluSx1 (ALA) is composed of an ERV1-encoding element in the middle and Alu elements at both ends (Figure 7D, Figure S15B). To further assess how ALA impacts cancer-specific attributes, we performed cell proliferation assay and scratch migration assay, finding that ALA-knockdown U251 cells showed significantly slower growth (Figure 7E) and migration (Figure 7F) relative to the control cells (Welch t-test, *p*-value < 0.05), supporting the reliability of TERA in detecting functional teRNAs.

## Discussion

While the majority of TEs are transcriptionally inactive, some have been domesticated to produce teRNAs that play crucial roles in a variety of biological processes and diseases. Thus, comprehensive benchmarking of teRNA analysis tools is in demand to reveal unbeknownst functional teRNAs, despite several challenges. The first challenge is the absence of a definitive reference for teRNAs. Although some full-length transcriptome datasets have been available for grabbing teRNA-transcripts, the low expression levels of teRNAs require much greater sequencing depth to establish a reliable ground truth for benchmarking. Secondly, the types of teRNAs are complex beyond TE units. Although traditionally assumed to arise from individual TE units, accumulating evidence demonstrated that TE units could be co-transcribed with neighboring genes and other TE units, producing TE-gene chimeric transcripts (chimeric teRNAs) or tandem-TE transcripts (a subclass of TE-only teRNAs). Our findings showed that less than 1% teRNA transcripts originate from a single TE unit without any genic sequence (Figure S2A), highlighting the need for benchmark teRNA identification beyond teRNA-units. Lastly, teRNAs have been analyzed across different levels (including family-, unit-, exon-, and transcript-levels), but yielded inconsistent results among levels due to algorithmic biases, calling for detailed comparison between algorithms.

With 120 simulated datasets generated from ultra-deep full-length transcriptome via the PacBio platform, we benchmarked the performance of 16 computational methods for teRNA detection and quantification across family-, unit-, exon-, and transcript-levels. We found that increased read length and sequencing depth contributed to better performance of teRNA detection in most software, especially for SERVE, but teRNA quantification remained unaffected. In terms of algorithm, reference-based assembly tools, particularly StringTie, were more efficient in detecting TE-gene chimeric teRNAs, while the *de novo* assembly method SERVE was the most sensitive for TE-only teRNAs. For teRNA quantification, Telescope showed outstanding performance at the unit-level and family-level, while RSEM and Kallisto demonstrated superior results at the transcript-level and exon-level.

A key finding of our study is the exceptional difficulty in accurately reconstructing full-length teRNA transcripts from short-read data, as evidenced by the low sensitivity of all tools. This challenge stems from several inherent limitations of short-read sequencing: (1) the limited read length rarely spans multiple exons, making splice graph resolution ambiguous; (2) the ubiquity of multi-mapping caused by high TE sequence similarity makes it intractable to accurately assign reads to specific transcript isoforms; and (3) the diversity and typically low expression abundance of TE transcripts lead to fragmented and incomplete assemblies. Consequently, even our best-practice pipeline, TERA, struggles with transcript-level identification. This fundamental limitation underscores the potential of long-read sequencing technologies (e.g., PacBio HiFi), which capture full-length transcripts and could dramatically improve teRNA identification. However, their current lower throughput presents a challenge for detecting low-abundance teRNAs and necessitates future independent benchmarking against experimental validation. Beyond technological advances, strategies such as targeted enrichment of TE-derived sequences could also revolutionize the field by making rare teRNAs detectable.

Following this benchmarking, we presented practical guidance for detecting and quantifying teRNAs from short-read RNA-seq data and developed the best-practice computational pipeline, TERA. TERA integrates existing teRNA analysis methods that showed good performance in teRNA detection and quantification at the family-, unit- or exon-levels, respectively. The selection of tools for TERA was guided by our benchmarking results: StringTie and SERVE were chosen for detection due to their complementary, high-performance, and computational efficiency in identifying chimeric and TE-only transcripts, respectively. For quantification, Telescope was selected for its superior accuracy at the unit and family levels, while RSEM and Kallisto were included as top-performing, practical options for the exon and transcript levels, offering users both alignment-based robustness and alignment-free speed. At the family-level, TERA surpassed existing tools in RNA-seq datasets with read length ≥ PE150 or sequencing depth ≥ 50 million pairs. TERA also excelled at the unit-level across all datasets and outperformed in exon-level detection and quantification from RNA-seq datasets with read length ≥ PE75 or sequencing depth ≥ 50 million pairs. Thus, TERA, as a user-friendly best-practice pipeline, is capable of reanalyzing most of public RNA-seq datasets for deciphering functional teRNAs.

Due to their repetitive nature, teRNA functions were often identified at the family-level for higher accuracy, posing challenges in determining whether a specific teRNA within a certain family is causally involved or merely a byproduct. To overcome this limit, several studies turned to locus-specific identification at the unit-level or transcript-level. However, our study highlighted teRNA-exon as a superior alternative. Unlike teRNA-units confined to single TE units, teRNA-exons accommodate chimeric teRNAs and tandem-TE teRNAs, which occupy the majority of teRNAs. Besides, teRNA-exons also provide additional detailed information, such as TE-TSS, TE-SS and TE-TTS. Using TERA on RNA-seq datasets from glioblastoma, we demonstrated the prospect of teRNA-exon analysis in identifying functional teRNAs with oncogenic potential. Thus, as an intermediate level between teRNA-transcript and teRNA-unit, teRNA-exons offer a balance between accuracy and resolution, providing novel insights for identification and functional interpretation of teRNAs.

Overall, our study provides a comprehensive benchmarking of teRNA detection and quantification tools and establishes a best-practice pipeline for teRNA analysis. The pipeline enhances the accuracy and reliability of teRNA studies, facilitating further research into the roles of teRNAs in various biological processes and diseases.

## Materials and methods

### Sample preparation

All cell lines were grown in a humidified incubator with 95% CO_2_ at 37 °C. HEK293T and U251 cell lines were cultured in DMEM medium (Catalog No. C11995500BT, Gibco, Carlsbad, CA) supplemented with 10% fetal bovine serum (Catalog No. CTCC-002-001, CTCC, Jinhua, China) and 100 U ml^−1^ penicillin-streptomycin (Catalog No. 15140122, Gibco). HSF cell lines were cultured in DMEM medium supplemented with 15% fetal bovine serum and 100 U ml^−1^ penicillin-streptomycin. All cells were passaged at 70–90% confluency with 0.05% Trypsin-EDTA (Catalog No. 25300054, Gibco). RNA was isolated from U251 and HEK293T cell pellets containing 1–5 × 10^6^ cells according to the manufacturer’s instructions using the FastPure^®^ Cell/Tissue Total RNA Isolation Kit V2 (Catalog No. RC112, Vazyme, Nanjing, China). Then, total RNA was divided into two equal parts and sent for strand-specific sequencing with 150-bp paired-end reads on Illumina platform and HiFi sequencing on PacBio platform at Novogene Co., Ltd (Beijing, China).

### Process of PacBio long-read HiFi sequencing data

We generated 164.73 million HiFi subreads, totaling 487.25 Gb of data from HEK293T and 190.45 million HiFi subreads, totaling 655.46 Gb of data from U251. CCS/HiFi reads were generated from subreads using ccs (v6.3.0). To generate high-quality full-length non-chimeric (FLNC) transcripts, we used lima (v2.4.0) for demultiplexing, IsoSeq3 (v3.4.0) for removing chimeric reads and LoRDEC [35] (v0.9) for correcting error with short-read RNA-seq data. Reads were then aligned with Minimap2 [36] (v2.17-r941) to human telomere-to-telomere (T2T) reference genome [37] (T2T-CHM13 v2.0) and collapsed into non-redundant full-length isoforms with cDNA_Cupcake (v29.0.0). With TE annotation from Dfam [38] (v3.8), we extracted TE transcripts (teRNAs) from FLNC transcripts by BEDTools [39] (v2.27.1). To obtain high-quality full-length teRNAs, we set strict criteria as follows: (1) for multi-exon transcripts, the supporting HiFi reads should be ≥ 2 or the splicing junction should be supported by ≥ 1 junction read from short-read RNA-seq data or transcript annotation (UCSC GENCODEv35 CAT/Liftoff v2); (2) for mono-exon transcripts, supporting HiFi reads should be ≥ 2; (3) sequence length of TEs within teRNAs should be ≥ 20 bp.

### Annotation of teRNAs

Our study categorized teRNAs into TE-gene chimeric teRNAs and TE-only teRNAs. TE-gene chimeric teRNAs were detected by aligning teRNAs to reference gene annotation (UCSC GENCODEv35 CAT/Liftoff v2) using GffCompare [40] (v0.12.6). TE loci within teRNAs were annotated with TE annotation from Dfam database using BEDTools. We also considered TE-only teRNAs with ≥ 2 TEs as tandem-TE teRNAs. TE-gene chimeric teRNAs were further divided into annotated chimeric teRNAs and novel chimeric teRNAs, depending on whether TEs within teRNAs are included in annotated transcripts from reference transcript annotation. Based on the location of teRNAs to genes, TE-only teRNAs were further categorized into intergenic teRNAs, intronic teRNAs and antisense teRNAs.

### Data simulation

Based on high-quality full-length teRNAs, we generated synthetic datasets using Polyester [41] (v1.22.0). Firstly, we combined all high-quality teRNAs and FLNC non-TE transcripts as transcript annotation for simulation. Then, the expression level of all transcripts were calculated from short-read RNA-seq data with RSEM (v1.2.28). Subsequently, whole-transcriptome simulation with empirically derived fragment GC bias was carried out using Polyester. Additionally, to benchmark tool performance across a range of expression levels, we created four simulated datasets from the same underlying transcriptome. The LTE dataset represents the ground truth containing all teRNAs at their original simulated expression levels (FPKM > 0). The MTE1, MTE2, and HTE datasets were created by generating entirely new expression values constrained to specific minimum thresholds (MTE1: FPKM > 0.05; MTE2: FPKM > 0.1; HTE: FPKM > 1) through a stratified random sampling process based on the empirical distribution of expression levels in our ground truth data and randomly reassigning these values to all transcripts. 20 million stranded paired-end 150-bp reads were generated for LTE, MTE1, MTE2 and HTE dataset.

### Public RNA-seq data

For benchmarking teRNA identification methods, we collected samples with both long-read and short-read RNA-seq data from Genotype-Tissue Expression Project (GTEx) through dbGaP and the NCBI Sequence Read Archive (SRA) (PRJNA635275). For benchmarking teRNA quantification methods, we downloaded HSF RNA-seq data from the NCBI SRA database (PRJNA776713) and Fibroblast-HERV annotation from GitHub (https://github.com/janky-yz/SERVE). RNA-seq data from glioblastoma and normal brain tissues were downloaded from the NCBI SRA database (PRJNA613939). RNA-seq data from glioblastoma patients and corresponding clinical data were downloaded from TCGA through GDC Data Prortal (https://portal.gdc.cancer.gov) and CGGA (http://www.cgga.org.cn).

### TERA implementation and usage

The TERA pipeline is implemented in Python and R. Its input requires RNA-seq reads in FASTQ format, the reference genome (in FASTA format), its gene annotation (in GTF format), and a custom TE annotation in both BED and GTF formats (the latter can be generated from the former using the provided script TEbedtogtf.R). For simplified deployment and to ensure reproducibility, we provide both a Conda environment (environment.yml) and a Singularity definition file (TERA.def) that encapsulate all software dependencies. The computational resources required are stage-dependent. The detect module is the most computationally intensive, typically requiring ∼16 hours, 40 threads, and ∼100 GB RAM for a sample with ∼20 million PE150 reads. The anno module is highly efficient (< 5 minutes), and the quant module’s runtime depends on the chosen tool (Telescope: ∼25 min; RSEM: ∼2.5 hours; Kallisto: ∼5 min). Detailed step-by-step instructions, including preparation of reference files, full command-line examples, and descriptions of all output files, are provided in the README file within the TERA GitHub repository (https://github.com/janky-yz/TERA).

### The benchmark evaluation process

To ensure the objectivity of the benchmark study, every evaluated method was tested with default parameters. For fair comparison, we applied the same reference genome (T2T-CHM13 v2.0), reference gene annotation (UCSC GENCODEv35 CAT/Liftoff v2), and TE annotation (all TE loci annotated in Dfam database for teRNA identification and all ground-truth teRNAs for teRNA quantification) in all methods for performance evaluation.

For teRNA identification, we calculated precision, sensitivity and F_1_-score for each evaluated method. For teRNA-transcript, we considered true positives (TP) as the query transcripts having the same TEs, the same number of exons and matched internal junctions (absolute error within ± 5 bp) with the ground truth, whereas the 5’ start or 3’ end of the first and last exon can vary. For teRNA-exon, TP were the query exons having the same TEs and matched internal junctions (absolute error within ± 5 bp) with the ground truth, whereas the 5’ start or 3’ end of the first and last exon can vary.

For teRNA quantification, we calculated Pearson’s correlation coefficient and root mean square error (RMSE) for each evaluated method. Absolute expression level of teRNA-exon is transformed from teRNA-transcript by length normalization. For TEtranscripts and SalmonTE, the teRNA-unit identifier was used in place of the teRNA-family name in order to generate locus-specific estimates. For Telescope, teRNA-family expression profiles were computed from teRNA-unit profiles by summing expression across all locations within each terRNA-family.

Benchmarking on real-world datasets was performed using two complementary strategies: (1) For public datasets where paired short-read and long-read data were available from the same sample, the long-read data was used to generate a high-confidence set of full-length teRNAs, which served as the ground truth for evaluating tool performance on the short-read data. (2) To provide orthogonal biological validation, we performed experimental validation (RT-PCR followed by Sanger sequencing for detection; RT-qPCR for quantification) on a subset of computationally predicted teRNAs.

### Experimental validation

For teRNA identification, we applied all evaluated methods in in-house short-read RNA-seq data from U251 and HEK293T cell lines and selected randomly 10 TEAM-specific teRNA-exons (including 5 chimeric teRNA-exons and 5 TE-only teRNA-exons) in each sample. The specificity of primer pairs was validated by aligning primer sequences to all annotated TEs and transcripts in the database using BLAST. All primer sequences are available in Table S10. Total RNA was reverse transcribed into cDNA by using a HiScript^®^ III 1st Strand cDNA Synthesis Kit (+ gDNA wiper) (Catalog No. R312, Vazyme), amplified with polymerase chain reaction (PCR) using 2× Rapid Taq Master Mix (Catalog No. P222, Vazyme) and validated by Sanger sequencing.

For teRNA quantification, we applied all methods in short-read RNA-seq data from HSFs with Fibroblast-HERV annotation and randomly selected 5 HERVs for reverse transcription–quantitative polymerase chain reaction (RT–qPCR) validation in HSFs using ChamQ Universal SYBR qPCR Master Mix (Catalog No. Q711, Vazyme). The threshold cycle (Ct) values of the selected HERVs were normalized to the housekeeping gene *GAPDH*. The primer sequences are shown in Table S10.

### Gene and teRNA expression analysis in RNA-seq data

For gene expression analysis, analysis protocol with STAR [42] (v2.7.5c) and featureCounts [43] (v2.0.0) was performed. In brief, the sequencing reads were aligned to the reference genome using STAR, and featureCounts was used to quantify gene expression levels. For teRNA expression analysis, TERA was used in teRNA identification at “detect” mode and teRNA quantification in all levels at “quant” mode. The similarity of gene expression and teRNA expression across samples was evaluated using multidimensional scaling. Differential expression analysis was performed using EdgeR [44] (v3.28.1) with raw read counts as the input. The Gene Ontology enrichment analysis and pathway enrichment analysis were conducted by the clusterProfiler [45] package (v3.14.3). Survival analysis was performed by the survival package (v3.7.0).

### Knockdown assay

Transfection of siRNAs (10 nM) were performed using INTERFERin^®^ transfection reagent (Catalog No. 101000028, Polyplus, Strasbourg, France) following the manufacturer’s instructions. The sequences of the siRNAs used in the present study are listed Table S11. The day before transfection, we seeded U251 cell lines in 6-well plates to obtain 30–50% confluency at the time of transfection. 6 hours after transfection, the supernatant was replaced with fresh cell medium.

### Cell proliferation assay

The wild-type cells and teRNA-knockdown cells were seeded into 96-well plates, at the starting density of 2500 cells per well. Ten microliters of Cell Counting Kit-8 (Catalog No. C0038, Beyotime, Shanghai, China) solution were added to each well at 24 hours, 48 hours, 72 hours and 96 hours. After 2 hours of incubation in humidified incubator with 95% CO_2_ at 37 °C, we recorded optical density at 450 nm.

### In vitro scratch migration assay

The wild-type cells and teRNA-knockdown cells were seeded into 6-well plates and grown to 100% confluency. We made straight scratches in middle of the well using 200-µl pipette tips and gently washed the well with culture medium twice to remove free floating cells. Then, we imaged the scratch with microscope and measured the area of the scratch using ImageJ at the time of the scratch, 12 hours, 24 hours and 48 hours after the scratch.

## Data availability

The raw sequence data generated in this study have been deposited in the NCBI SRA database (BioProject: PRJNA1138880) and Genome Sequence Archive [46] at the National Genomics Data Center (NGDC), Beijing Institute of Genomics (BIG), Chinese Academy of Sciences (CAS) / China National Center for Bioinformation (CNCB) (BioProject: PRJCA047046), which are publicly accessible at https://ngdc.cncb.ac.cn/gsa. Details about data analyzed in this study were included in the Materials and methods section.

## Code availability

All of the scripts used to compute the metrics described in the study, source code for TERA and example data, are available at GitHub (https://github.com/janky-yz/TERA), as well as BioCode at the NGDC, BIG, CAS / CNCB (BioCode: BT008011), which is publicly accessible at https://ngdc.cncb.ac.cn/biocode/tools/8011.

## Author contributions

E.Y. conceived and designed the study. J.Q.S. and J.D.W. performed the experiments.

J.Q.S. analyzed the data. J.Q.S. and E.Y. interpreted the data and wrote the manuscript.

All authors approved the final version of the manuscript.

## Acknowledgments

The Genotype-Tissue Expression (GTEx) Project was supported by the Common Fund of the Office of the Director of the National Institutes of Health, and by NCI, NHGRI, NHLBI, NIDA, NIMH, and NINDS. This work was supported by grant from the Ministry of Science and Technology of China (2021ZD0203203), Young Scientists Fund of the National Natural Science Foundation of China (32400534) and the China Postdoctoral Science Foundation (2024M760136).

## References

[1] International Human Genome Sequencing Consortium. Initial sequencing and analysis of the human genome. Nature 2001;409:860–921.

[2] Chenais B, Caruso A, Hiard S, Casse N. The impact of transposable elements on eukaryotic genomes: from genome size increase to genetic adaptation to stressful environments. Gene 2012;509:7–15.

[3] Slotkin RK, Martienssen R. Transposable elements and the epigenetic regulation of the genome. Nat Rev Genet 2007;8:272–285.

[4] Orgel LE, Crick FH. Selfish DNA: the ultimate parasite. Nature 1980;284:604–607.

[5] Suntsova M, Garazha A, Ivanova A, Kaminsky D, Zhavoronkov A, Buzdin A. Molecular functions of human endogenous retroviruses in health and disease. Cell Mol Life Sci 2015;72:3653–3675.

[6] Zhang Y, Li T, Preissl S, Amaral ML, Grinstein JD, Farah EN, et al. Transcriptionally active HERV-H retrotransposons demarcate topologically associating domains in human pluripotent stem cells. Nat Genet 2019;51:1380–1388.

[7] Zhou X, Singh M, Santos GS, Guerlavais V, Carvajal LA, Aivado M, et al. Pharmacologic activation of p53 triggers viral mimicry response thereby abolishing tumor immune evasion and promoting antitumor immunity. Cancer Discov 2021;11:3090–3105.

[8] Liu X, Liu Z, Wu Z, Ren J, Fan Y, Sun L, et al. Resurrection of endogenous retroviruses during aging reinforces senescence. Cell 2023;186:287–304.

[9] Jang HS, Shah NM, Du AY, Dailey ZZ, Pehrsson EC, Godoy PM, et al. Transposable elements drive widespread expression of oncogenes in human cancers. Nat Genet 2019;51:611–617.

[10] Shah NM, Jang HJ, Liang Y, Maeng JH, Tzeng SC, Wu A, et al. Pan-cancer analysis identifies tumor-specific antigens derived from transposable elements. Nat Genet 2023;55:631–639.

[11] Scopa C, Barnada SM, Cicardi ME, Singer M, Trotti D, Trizzino M. JUN upregulation drives aberrant transposable element mobilization, associated innate immune response, and impaired neurogenesis in Alzheimer’s disease. Nat Commun 2023;14:8021.

[12] Lu X, Sachs F, Ramsay L, Jacques PÉ, Göke J, Bourque G, et al. The retrovirus HERVH is a long noncoding RNA required for human embryonic stem cell identity. Nat Struct Mol Biol 2014;21:423–425.

[13] Ng KW, Boumelha J, Enfield KSS, Almagro J, Cha H, Pich O, et al. Antibodies against endogenous retroviruses promote lung cancer immunotherapy. Nature 2023;616:563–573.

[14] Lanciano S, Cristofari G. Measuring and interpreting transposable element expression. Nat Rev Genet 2020;21:721–736.

[15] Iñiguez LP, de Mulder Rougvie M, Stearrett N, Jones RB, Ormsby CE, Reyes-Terán G, et al. Transcriptomic analysis of human endogenous retroviruses in systemic lupus erythematosus. Proc Natl Acad Sci U S A 2019;116:21350–21351.

[16] Pertea M, Pertea GM, Antonescu CM, Chang TC, Mendell JT, Salzberg SL. StringTie enables improved reconstruction of a transcriptome from RNA-seq reads. Nat Biotechnol 2015;33:290–295.

[17] Trapnell C, Williams BA, Pertea G, Mortazavi A, Kwan G, van Baren MJ, et al. Transcript assembly and quantification by RNA-Seq reveals unannotated transcripts and isoform switching during cell differentiation. Nature Biotechnol 2010;28:511–515.

[18] Babaian A, Thompson IR, Lever J, Gagnier L, Karimi MM, Mager DL. LIONS: analysis suite for detecting and quantifying transposable element initiated transcription from RNA-seq. Bioinformatics 2019;35:3839–3841.

[19] She J, Du M, Xu Z, Jin Y, Li Y, Zhang D, et al. The landscape of hervRNAs transcribed from human endogenous retroviruses across human body sites. Genome Biol 2022;23:231.

[20] Bendall ML, de Mulder M, Iñiguez LP, Lecanda-Sánchez A, Pérez-Losada M, Ostrowski MA, et al. Telescope: characterization of the retrotranscriptome by accurate estimation of transposable element expression. PLoS Comput Biol 2019;15:e1006453.

[21] Jin Y, Tam OH, Paniagua E, Hammell M. TEtranscripts: a package for including transposable elements in differential expression analysis of RNA-seq datasets. Bioinformatics 2015;31:3593–3599.

[22] Lerat E, Fablet M, Modolo L, Lopez-Maestre H, Vieira C. TE_TOOLS_ facilitates big data expression analysis of transposable elements and reveals an antagonism between their activity and that of piRNA genes. Nucleic Acids Res 2017;45:e17.

[23] Yang WR, Ardeljan D, Pacyna CN, Payer LM, Burns KH. SQuIRE reveals locus-specific regulation of interspersed repeat expression. Nucleic Acids Res 2019;47:e27.

[24] Jeong HH, Yalamanchili HK, Guo C, Shulman JM, Liu Z. An ultra-fast and scalable quantification pipeline for transposable elements from next generation sequencing data. Pac Symp Biocomput 2018;23:168–179.

[25] Criscione SW, Zhang Y, Thompson W, Sedivy JM, Neretti N. Transcriptional landscape of repetitive elements in normal and cancer human cells. BMC genomics 2014;15:583.

[26] Li B, Dewey CN. RSEM: accurate transcript quantification from RNA-Seq data with or without a reference genome. BMC Bioinformatics 2011;12:323.

[27] Roberts A, Pachter L. Streaming fragment assignment for real-time analysis of sequencing experiments. Nat Methods 2013;10:71–73.

[28] Patro R, Duggal G, Love MI, Irizarry RA, Kingsford C. Salmon provides fast and bias-aware quantification of transcript expression. Nat Methods 2017;14:417–419.

[29] Bray NL, Pimentel H, Melsted P, Pachter L. Near-optimal probabilistic RNA-seq quantification. Nat Biotechnol 2016;34:525–527.

[30] The GTEx Consortium. The GTEx Consortium atlas of genetic regulatory effects across human tissues. Science 2020;369:1318–1330.

[31] Huang KK, Huang J, Wu JKL, Lee M, Tay ST, Kumar V, et al. Long-read transcriptome sequencing reveals abundant promoter diversity in distinct molecular subtypes of gastric cancer. Genome Biol 2021;22:44.

[32] Huang T, Yang Y, Song X, Wan X, Wu B, Sastry N, et al. PRMT6 methylation of RCC1 regulates mitosis, tumorigenicity, and radiation response of glioblastoma stem cells. Mol Cell 2021;81:1276–1291.e9.

[33] Brennan CW, Verhaak RG, McKenna A, Campos B, Noushmehr H, Salama SR, et al. The somatic genomic landscape of glioblastoma. Cell 2013;155:462–477.

[34] Zhao Z, Zhang KN, Wang Q, Li G, Zeng F, Zhang Y, et al. Chinese Glioma Genome Atlas (CGGA): A comprehensive resource with functional genomic data from Chinese glioma patients. Genomics Proteomics Bioinformatics 2021;19:1–12.

[35] Salmela L, Rivals E. LoRDEC: accurate and efficient long read error correction. Bioinformatics 2014;30:3506–3514.

[36] Li H. Minimap2: pairwise alignment for nucleotide sequences. Bioinformatics 2018;34:3094–3100.

[37] Nurk S, Koren S, Rhie A, Rautiainen M, Bzikadze AV, Mikheenko A, et al. The complete sequence of a human genome. Science 2022;376:44–53.

[38] Storer J, Hubley R, Rosen J, Wheeler TJ, Smit AF. The Dfam community resource of transposable element families, sequence models, and genome annotations. Mob DNA 2021;12:2.

[39] Quinlan AR, Hall IM. BEDTools: a flexible suite of utilities for comparing genomic features. Bioinformatics 2010;26:841–842.

[40] Pertea G, Pertea M. GFF Utilities: GffRead and GffCompare. F1000Res 2020;9:304.

[41] Frazee AC, Jaffe AE, Langmead B, Leek JT. Polyester: simulating RNA-seq datasets with differential transcript expression. Bioinformatics 2015;31:2778–2784.

[42] Dobin A, Davis CA, Schlesinger F, Drenkow J, Zaleski C, Jha S, et al. STAR: ultrafast universal RNA-seq aligner. Bioinformatics 2013;29:15–21.

[43] Liao Y, Smyth GK, Shi W. featureCounts: an efficient general purpose program for assigning sequence reads to genomic features. Bioinformatics 2014;30:923–930.

[44] Robinson MD, McCarthy DJ, Smyth GK. edgeR: a Bioconductor package for differential expression analysis of digital gene expression data. Bioinformatics 2010;26:139–140.

[45] Yu G, Wang LG, Han Y, He QY. clusterProfiler: an R package for comparing biological themes among gene clusters. OMICS 2012;16:284–287.

[46] Chen T, Chen X, Zhang S, Zhu J, Tang B, Wang A, et al. The Genome Sequence Archive Family: toward explosive data growth and diverse data types. Genomics Proteomics Bioinformatics 2021;19:578–583.

