## Supplementary Figures 1-15 for "Comprehensive benchmarking with guidelines for analyzing transposable element-derived RNA expression"

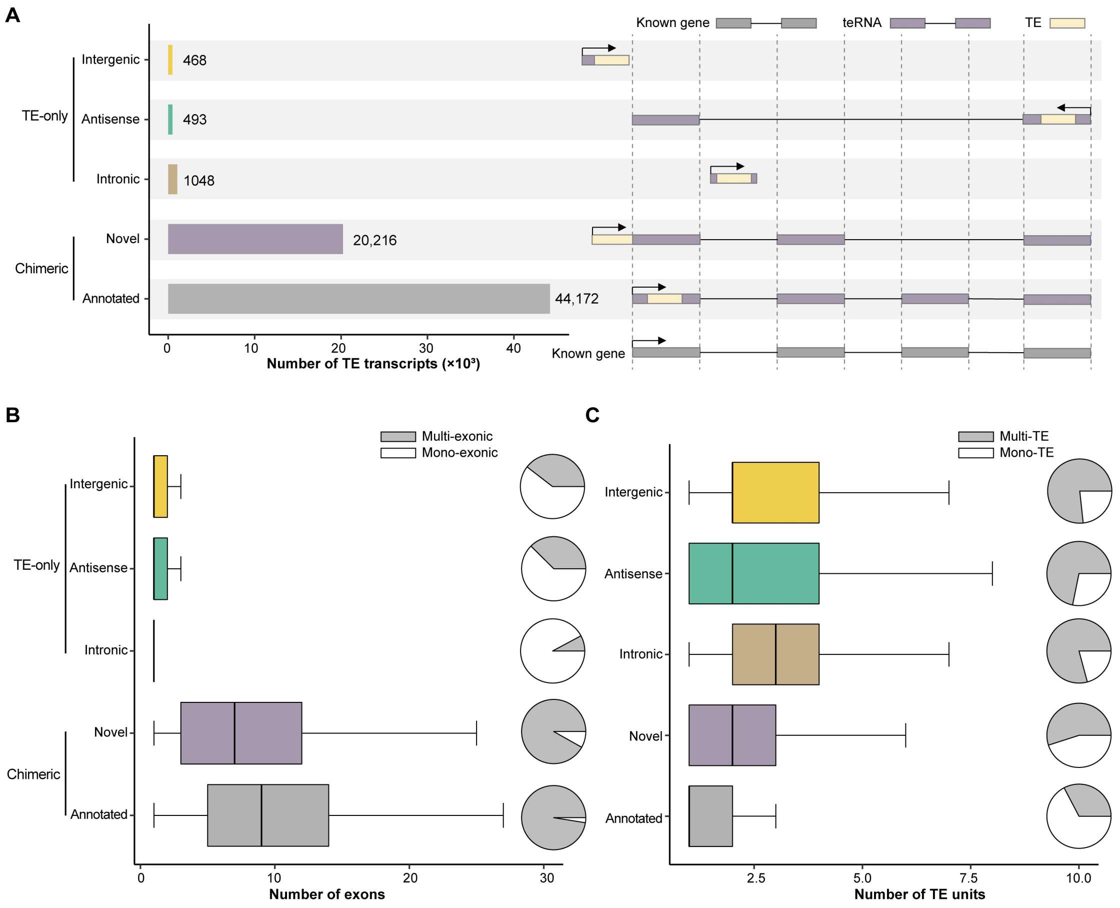


**Figure S1 Full-length teRNAs in HEK293T cells.**

**A**. The number of full-length chimeric teRNAs (annotated and novel) and TE-only teRNAs (intergenic, antisense and intronic) detected from HEK293T cells. **B**–**C**. The number of exons (**B**) and TE units (**C**) of teRNAs stratified by their transcript classification in HEK293T cells. The center line indicates the median, the limits are the interquartile range (IQR), the whiskers represent 1.5× the IQR.

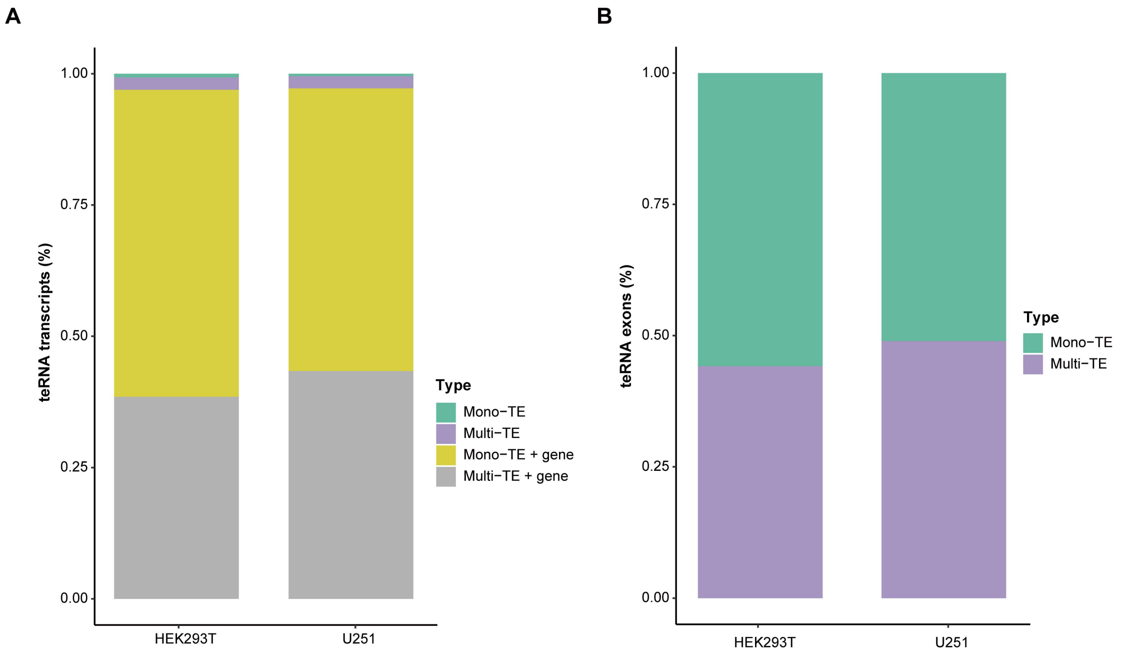


**Figure S2 The distribution of different types of teRNAs.**

**A**. The proportion of different types of teRNA-transcripts. **B**. The proportion of multi-TE and mono-TE exons.

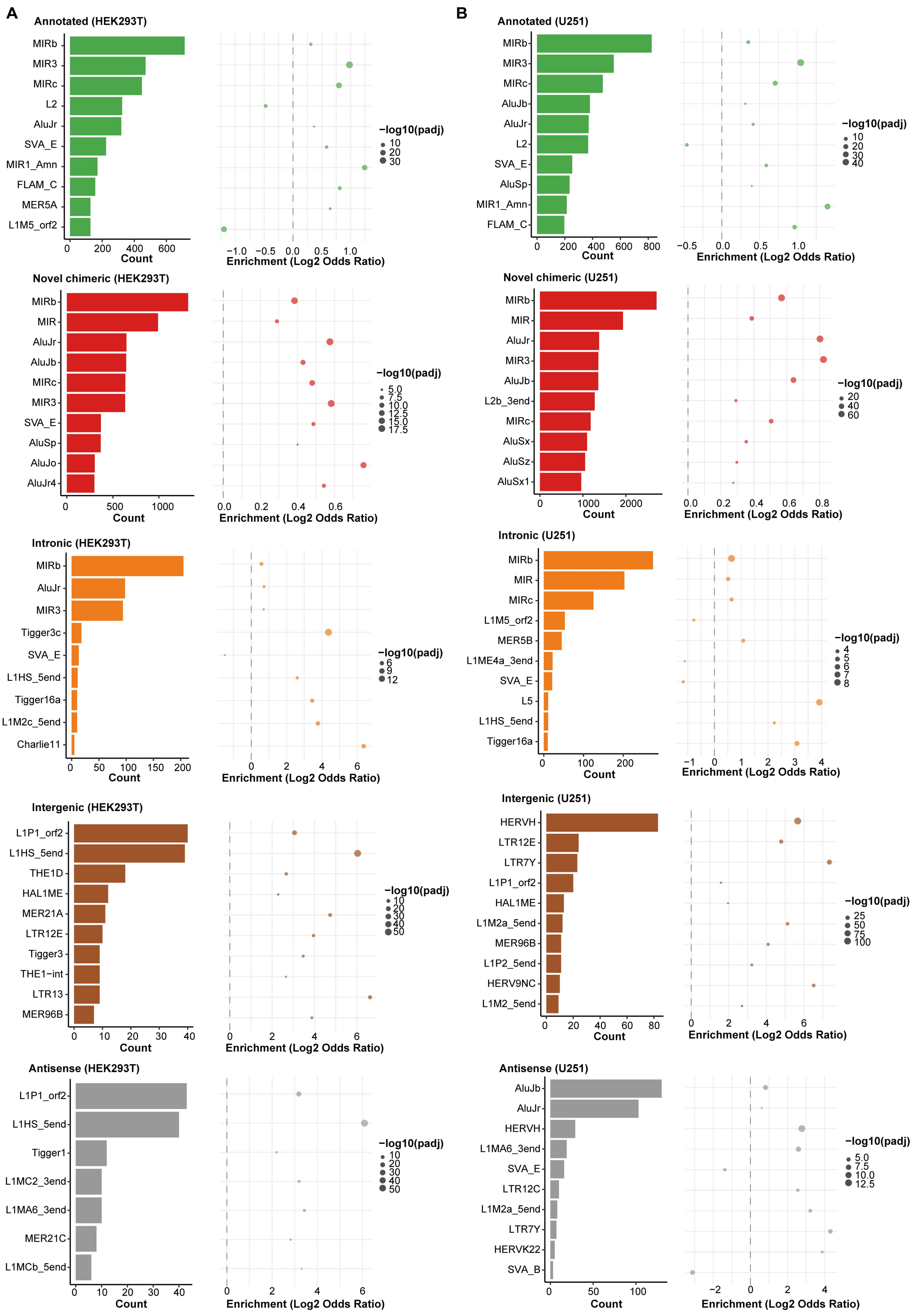


**Figure S3 TE family enrichment analysis on TE exons.**

**A–B**. The prevalence and enrichment of TE families on different types of TE exons in HEK293T (**A**) and U251 (**B**) cell lines.

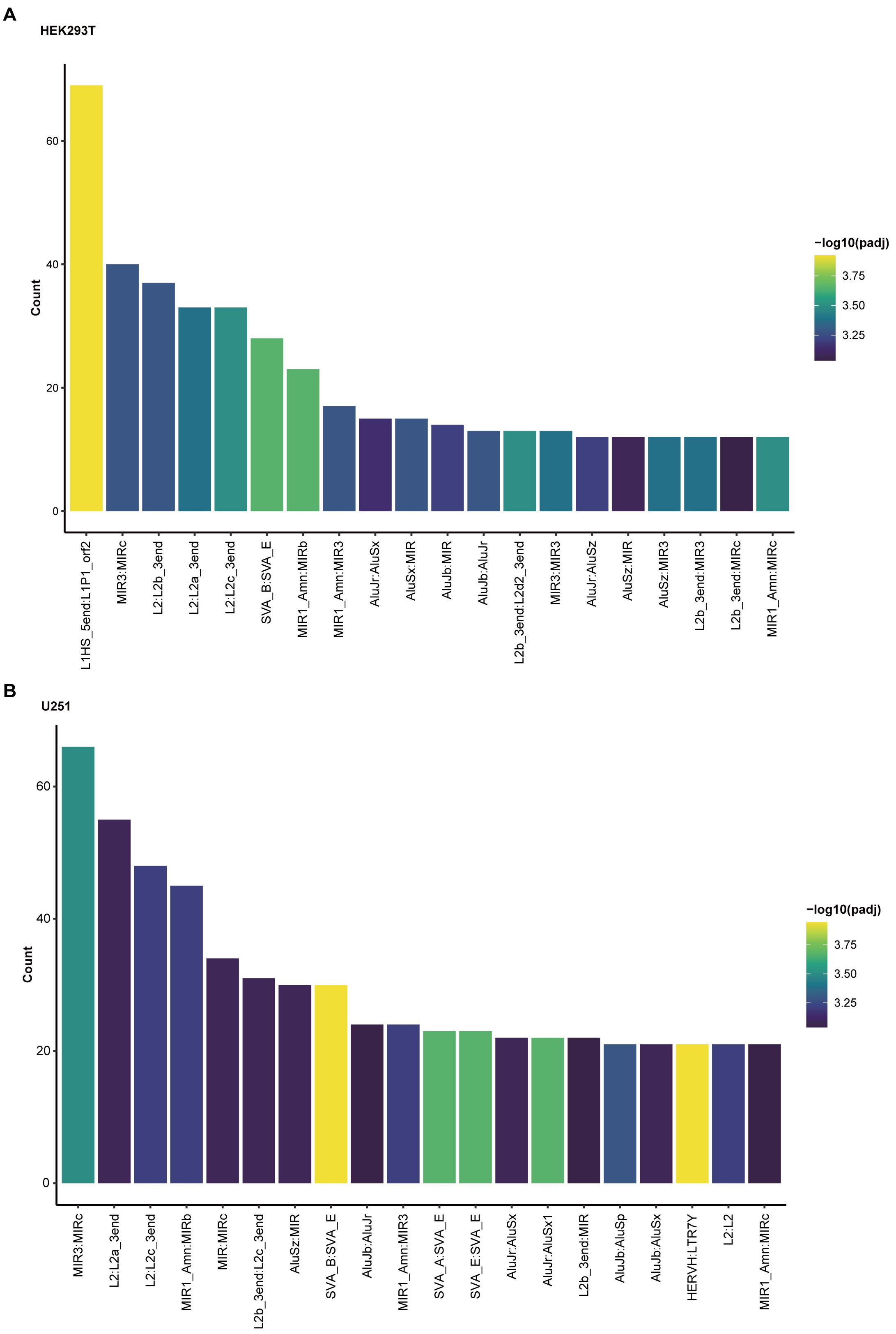


**Figure S4 The prevalence of TE unit co-occurrence patterns within TE exons.**

**A–B**. The prevalence and enrichment of TE combinations within exons in HEK293T (**A**) and U251 (**B**) cell lines.

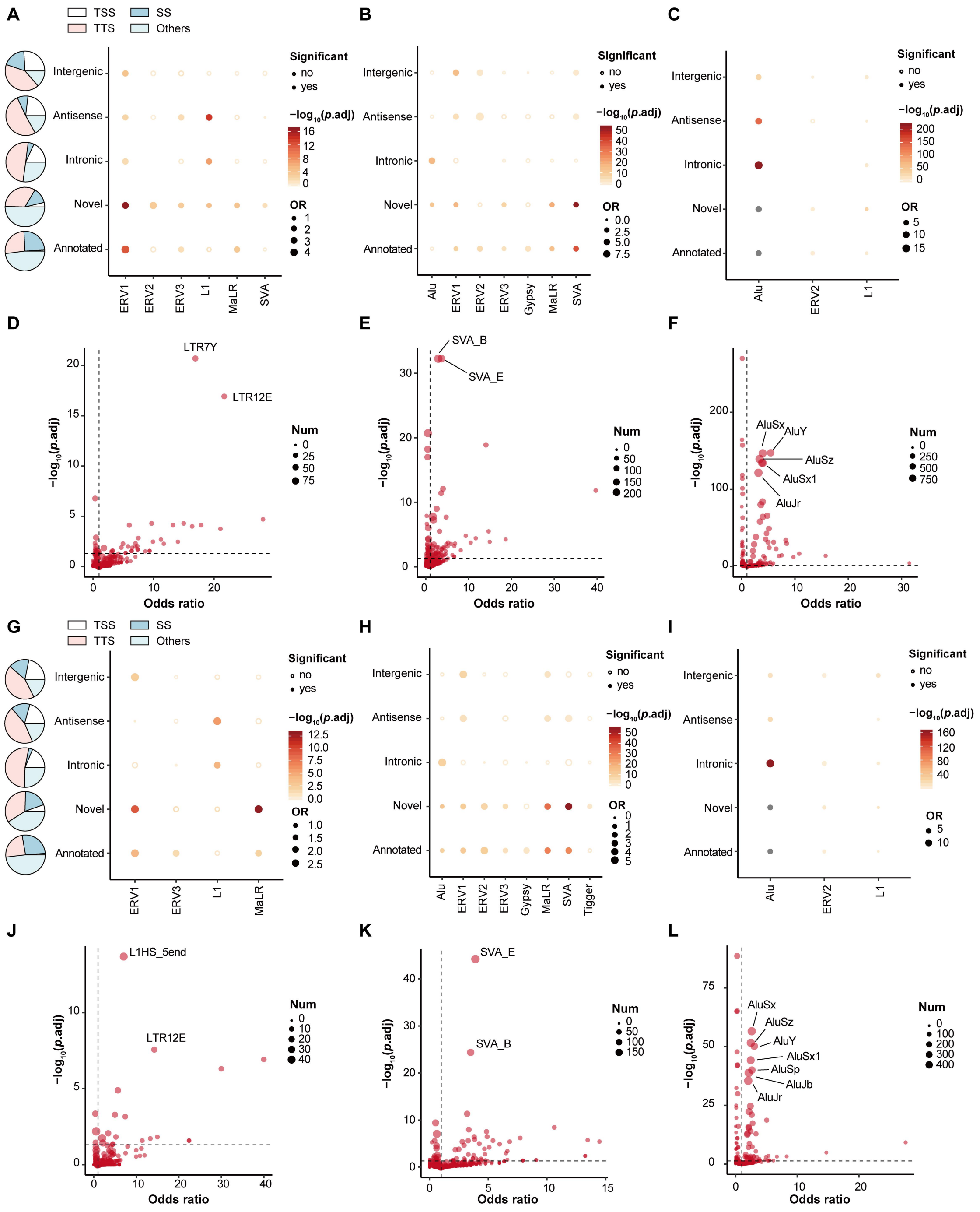

**Figure S5 TE family enrichment analysis on teRNAs.**

**A–C**. TE family enrichment analysis on TE-TSS teRNAs (**A**), TE-SS teRNAs (**B**) and TE-TTS teRNAs (**C**) in U251 cells. **D–F**. TE subfamily enrichment analysis on TE-TSS teRNAs (**D**), TE-SS teRNAs (**E**) and TE-TTS teRNAs (**F**) in U251 cells. **G–I**. TE family enrichment analysis on TE-TSS teRNAs (**G**), TE-SS teRNAs (**H**) and TE-TTS teRNAs (**I**) in

HEK293T cells. **J-L**. TE subfamily enrichment analysis on TE-TSS teRNAs (**J**), TE-SS teRNAs (**K**) and TE-TTS teRNAs (**L**) in HEK293T cells.

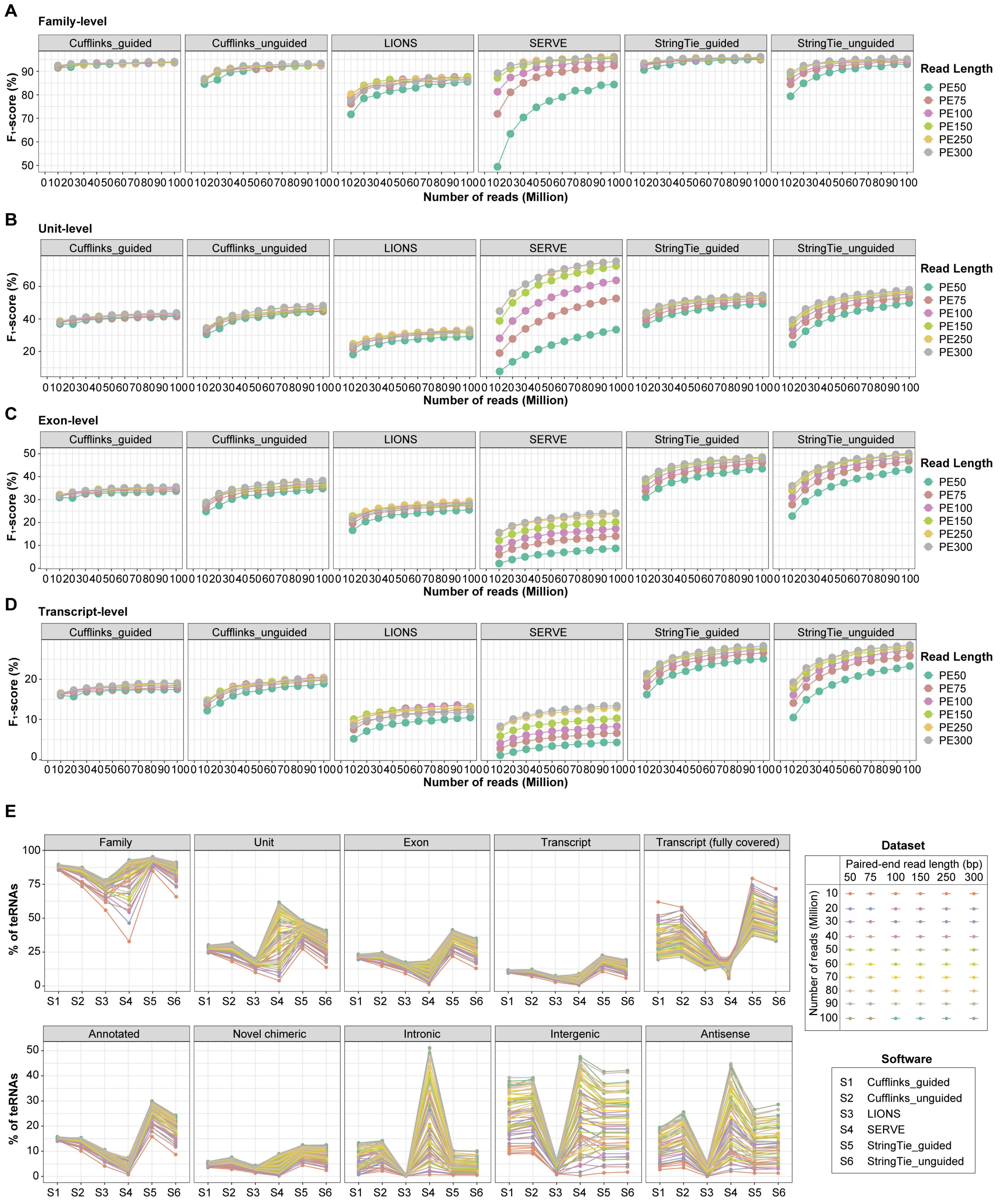


**Figure S6 Evaluation of teRNA detection methods across different levels.**

**A–D**. The performance of teRNA detection methods at the family- (**A**), unit- (**B**), exon- (**C**) and transcript-levels (**D**) in simulated datasets from HEK293T cells. **E**. Sensitivity of teRNA detection methods in the identification of different levels and types of teRNAs in simulated datasets from HEK293T cells. The center line indicates the median, the limits are IQR, the whiskers represent 1.5× the IQR.

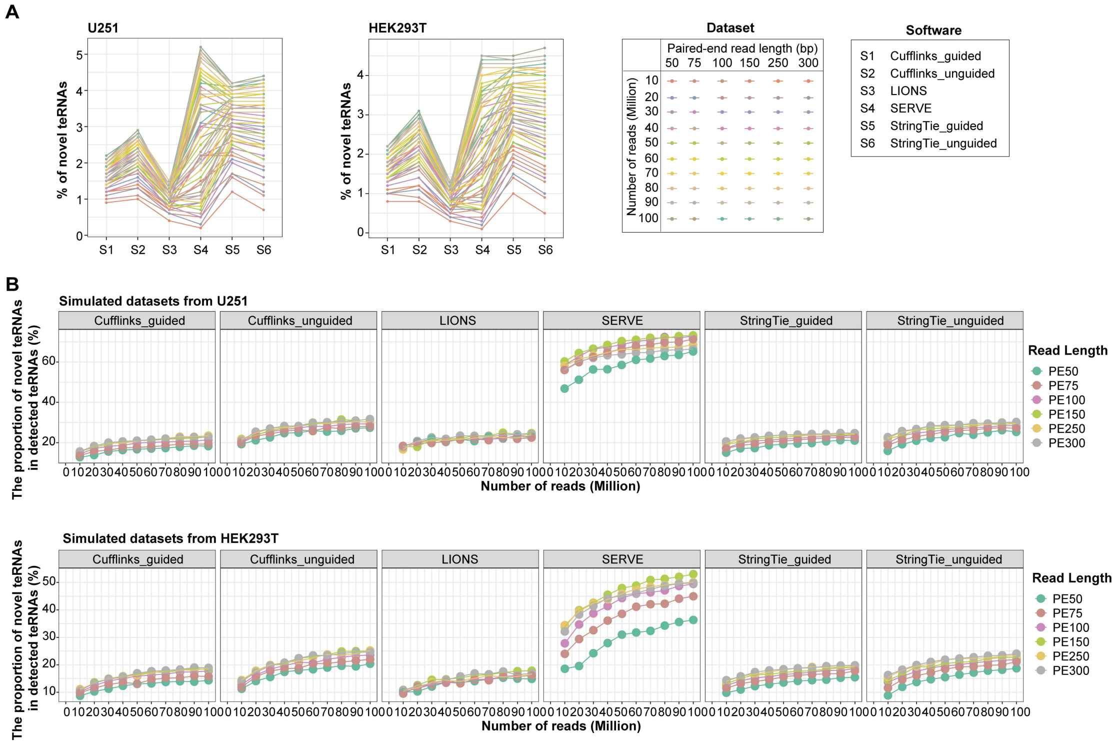


**Figure S7 Detection of novel teRNAs by different methods.**

**A**. Sensitivity of teRNA detection methods in the identification of novel teRNAs in simulated datasets. **B–C**. The proportion of novel teRNA in detected teRNAs across simulated datasets from U251 (**B**) and HEK293T (**C**) cells.

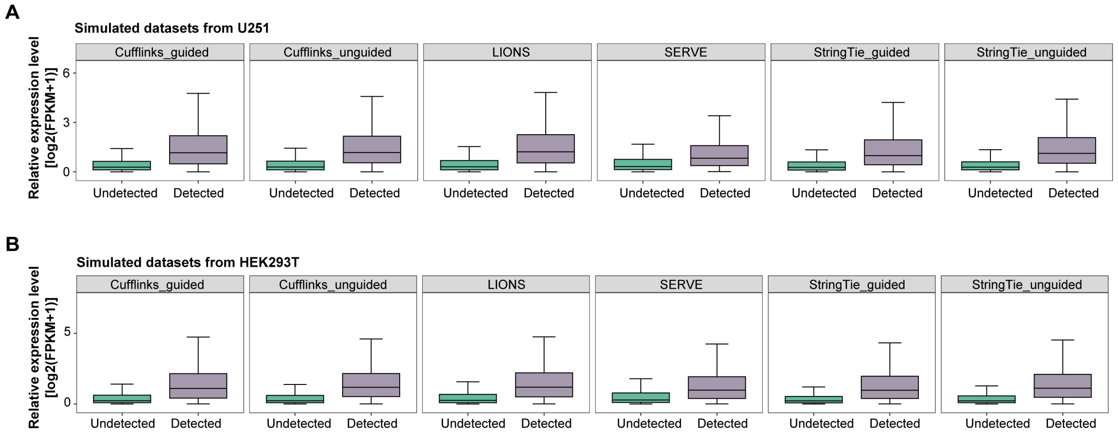


**Figure S8 The influence factors of teRNA detection.**

**A–B**. The expression of detected teRNA-transcripts showed significantly higher than undetected teRNA-transcripts in simulated datasets from U251 (**A**) and HEK293T (**B**) cells for all methods. The center line indicates the median, the limits are IQR, the whiskers represent 1.5× the IQR.

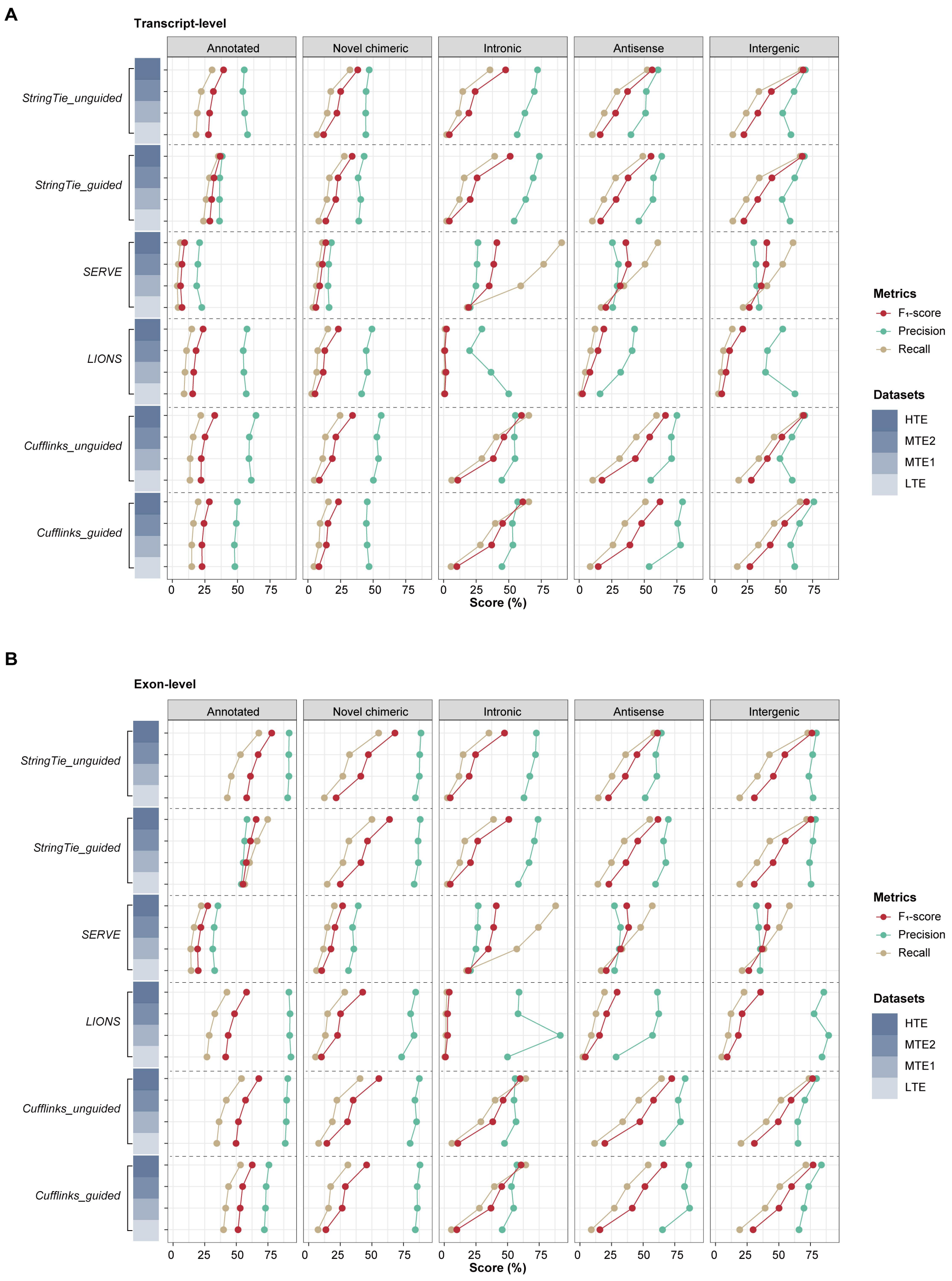


**Figure S9 Performance of methods in the detection of teRNAs with different expression level.**

**A–B**. The performance of teRNA detection methods in the identification of different types of teRNA-transcripts (**A**) teRNA-exons (**B**) in LTE, MTE1, MTE2 and HTE datasets from HEK293T cells.

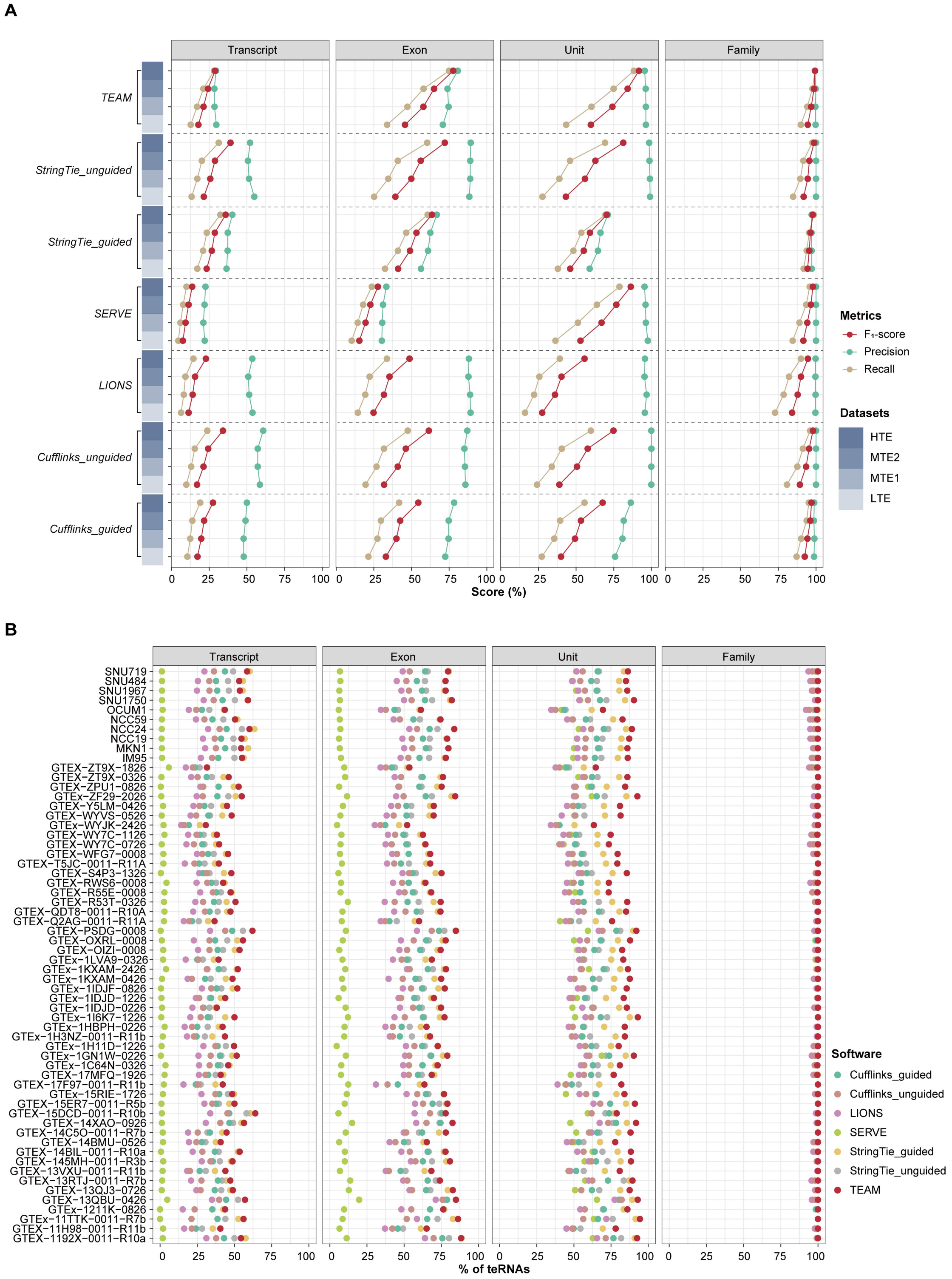


**Figure S10 The performance of TEAM and other teRNA detection methods.**

**A**. The performance of TEAM and existing teRNA detection methods across different levels in benchmarking simulated datasets from HEK293T cells. **B**. The sensitivity of teRNA detection methods in the identification of teRNAs across different levels in real-world datasets.

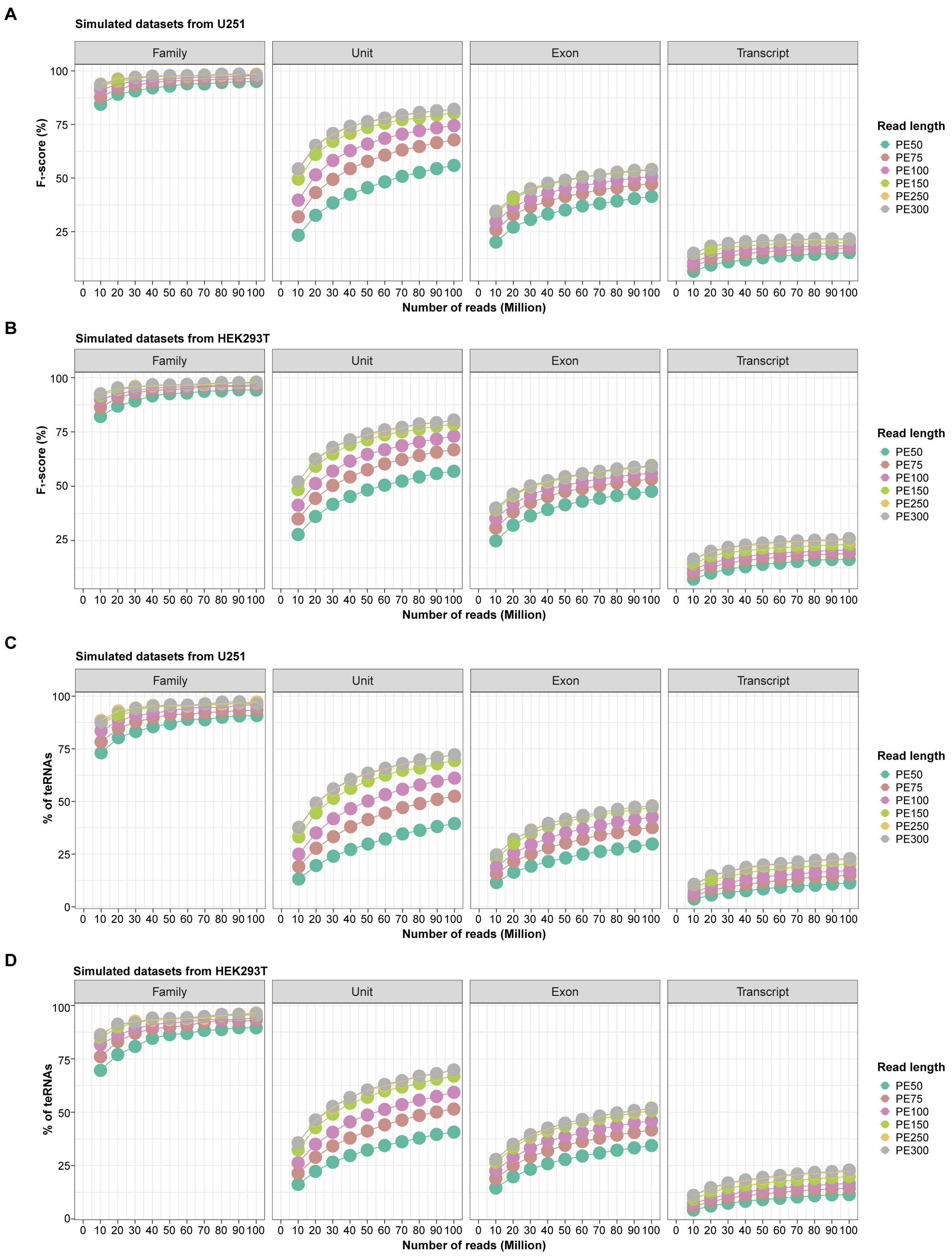


**Figure S11 Evaluation of TEAM on teRNA detection.**

**A–B**. The performance of TEAM across different levels in benchmarking simulated datasets from U251 (**A**) and HEK293T (**B**) cells. **C–D**. The sensitivity of TEAM across different levels in benchmarking simulated datasets from U251 (**A**) and HEK293T (**B**) cells.

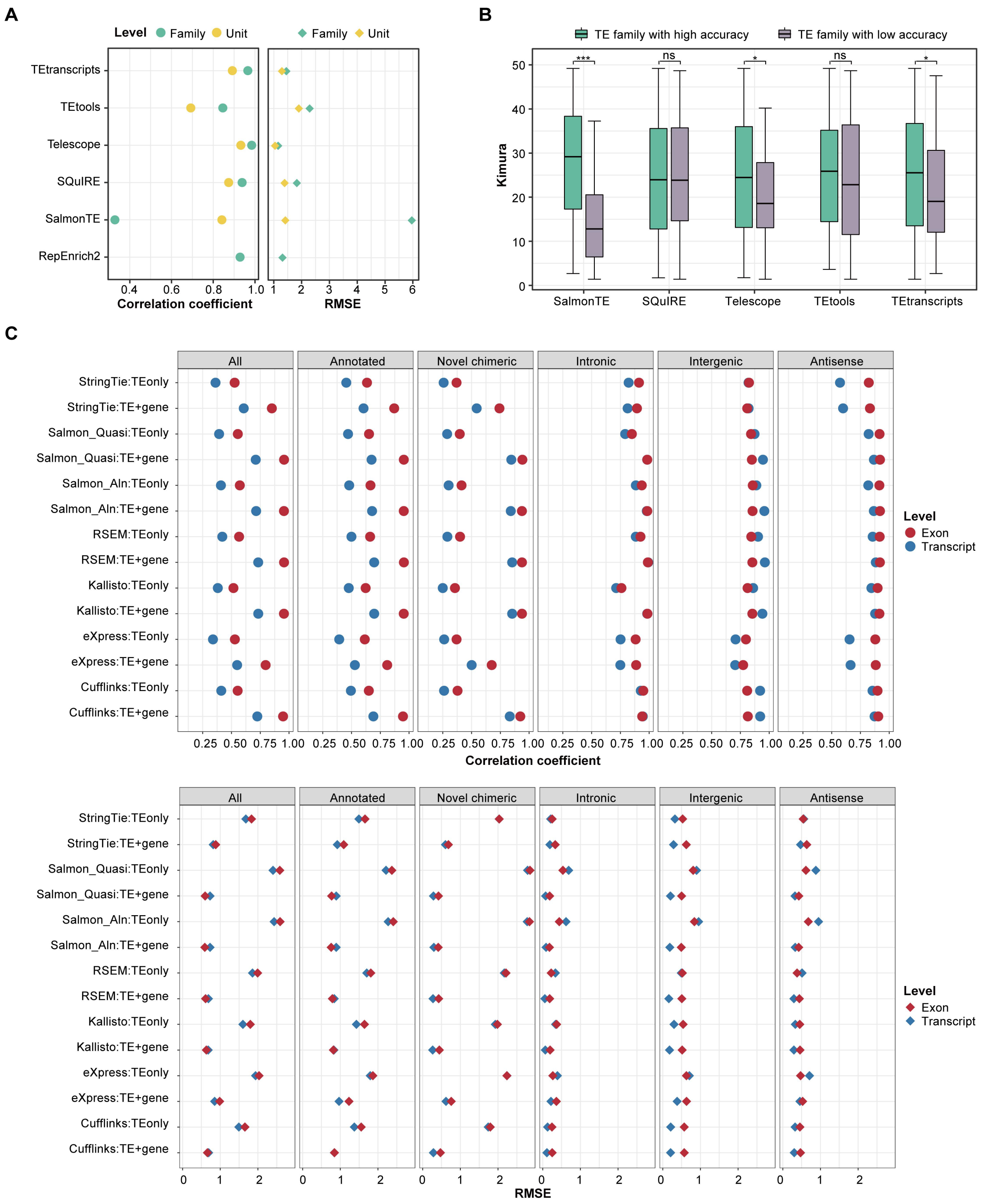


**Figure S12 Evaluation of teRNA quantification methods in simulated datasets from HEK293T cells.**

**A**. The performance of teRNA quantification methods at the teRNA-unit and teRNA-family level. **B**. Kimura's distance of teRNA-families with high and low quantification accuracy in HEK293T cells. The center line indicates the median, the limits are IQR, the whiskers represent 1.5× the IQR. *, $p$-value < 0.05; **, $p$-value < 0.01; ***, $p$-value < 0.001. **C**. The performance of teRNA quantification methods at the transcript-level and exon-level.

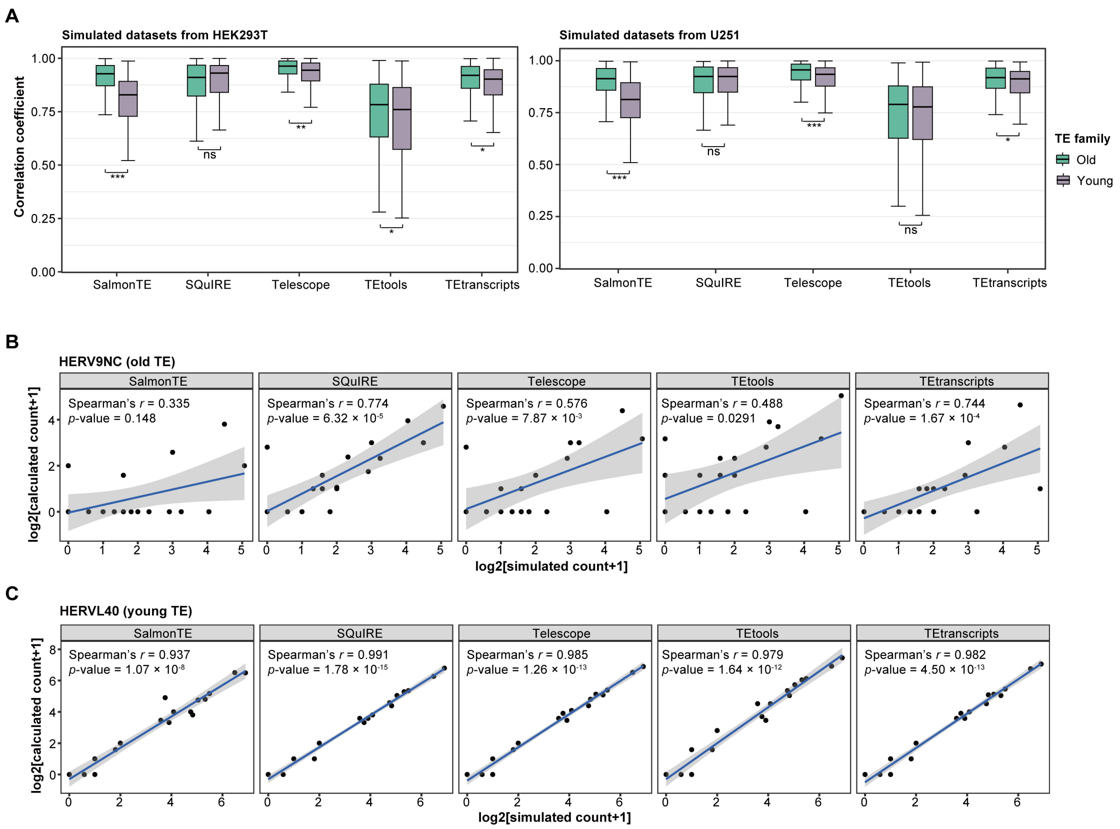


**Figure S13 Quantification accuracy of young and old TE families by different methods.**

**A**. The performance of teRNA quantification methods at the teRNA-unit level for young and old TE families. **B–C**. The performance of teRNA quantification methods on HERV9NC (**B**) and HERVL40 (**C**) family.

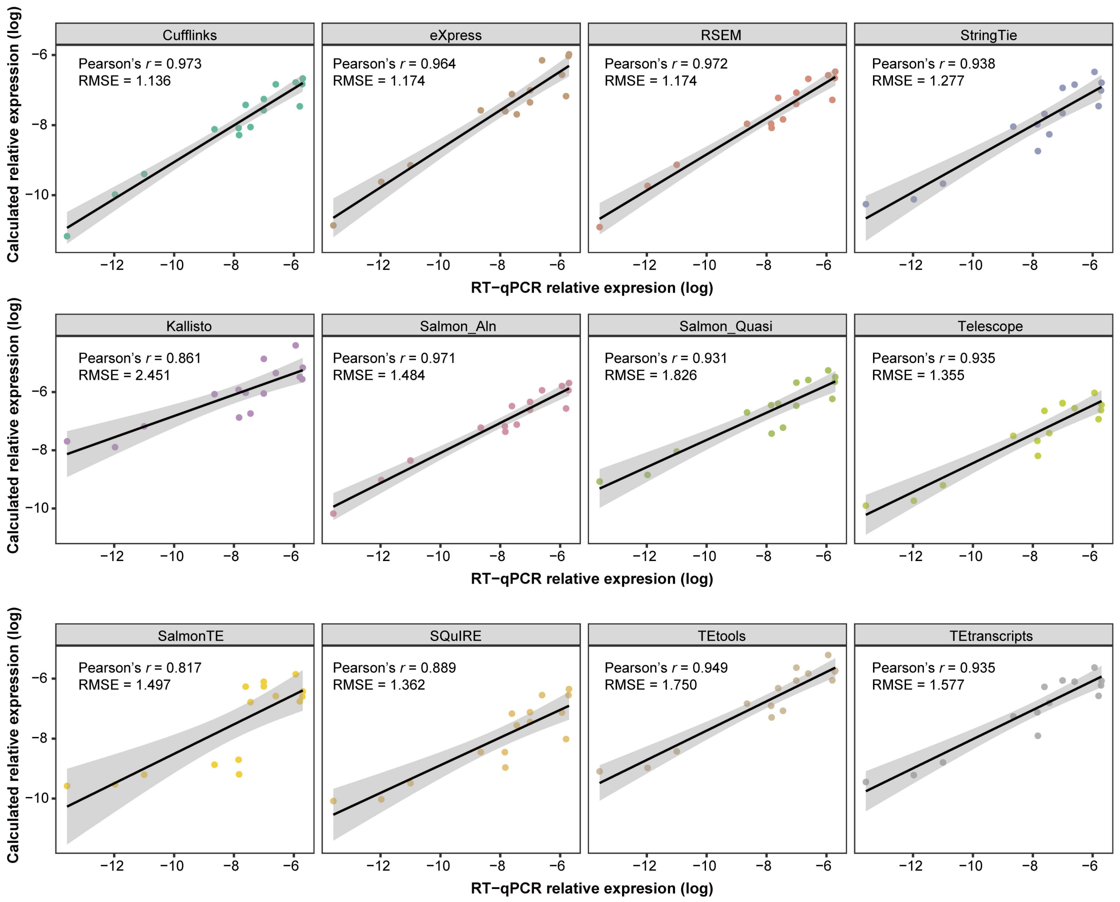


**Figure S14 Experimental validation of teRNA quantification methods. The performance of all teRNA quantification methods in HSF cells, measured by RT-qPCR.**

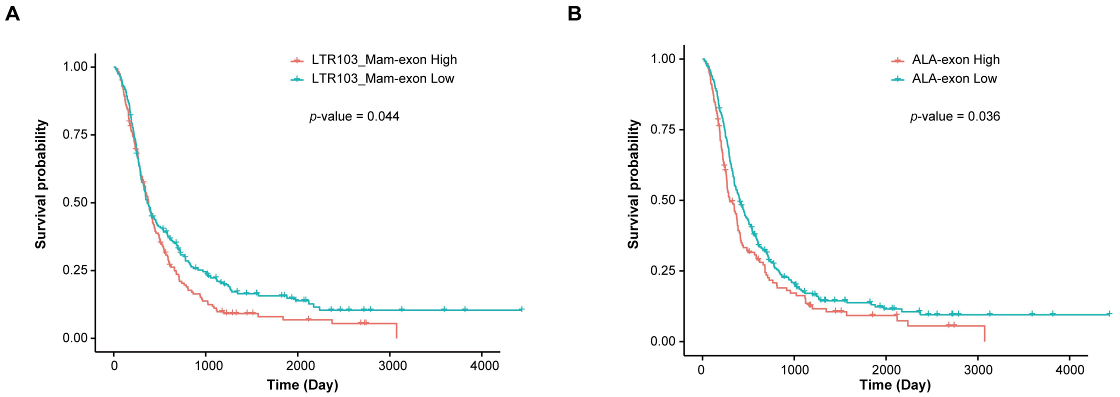


**Figure S15 Kaplan-Meier analysis of overall survival in glioblastoma.**

**A–B**. Kaplan-Meier analysis of overall survival showed that LTR103_Mam-exon (**A**) and ALA-exon (**B**) expression was significantly associated with worse prognosis in glioblastoma patients from CGGA datasets.
